# Molecular evidence for pre-chordate origins of ovarian cell types and neuroendocrine control of reproduction

**DOI:** 10.1101/2025.03.24.644836

**Authors:** Periklis Paganos, Carsten Wolff, Danila Voronov, S. Zachary Swartz

## Abstract

Sexual reproduction in animals requires the development of oocytes, or egg cells. This process, termed oogenesis, requires complex interactions amongst germline and somatic cell types in the ovary. How did these cell types and their signaling interactions evolve? Here we use the sea star *Patiria miniata* as a non-chordate deuterostome representative to define the ovarian cell type toolkit in echinoderms. Sea stars continuously produce millions of new oocytes throughout their lifespan, making them a practical system to understand the mechanisms that drive oogenesis from a biomedical and evolutionary perspective. We performed scRNA-seq combined with high-resolution 3D-imaging to reveal the ovarian cell types and their spatial organization. Our data support the presence of actively dividing oogonial stem cells and granulosa-like and theca-like cells, which display similarities and possible homology with their mammalian counterparts. Lastly, our data support the existence of an endocrine signaling system between oogonial stem cells and intrinsic ovarian neurons with striking similarities to the vertebrate hypothalamic-pituitary-gonadal axis. Overall, this study provides molecular evidence supporting the possible pre-chordate origins of conserved ovarian cell types, and the presence of an intrinsic neuroendocrine system which potentially controls oogenesis and predates the formation of the hypothalamic-pituitary-gonadal axis in vertebrates.

## Introduction

Oogenesis is essential for the propagation of most animal species. This process takes place within the ovary, where somatic and germ cells interact to form a fundamental reproductive unit: the follicle (1). Diverse ovarian cell types underlie these interactions and define the reproductive potential of the organism. For instance, somatic follicle cells, also referred to as granulosa in mammals, are in direct contact with developing oocytes and provide signals and metabolic support to regulate growth and meiotic arrest. Mammalian theca cells in the outer layers of the follicles have a key role in steroid hormone production, nutrient delivery to the follicle through the vasculature, and regulating granulosa cell function (2). In vertebrates, the hypothalamic-pituitary gland-gonadal (HPG) axis provides hormonal cues from the central nervous system essential for proper germ cell growth, maturation, and ovulation (3).

Many germ line processes are broadly conserved, but different organisms have diverse reproductive potentials due to the presence or absence of specific cell types or states. For instance, the mammalian oocyte reserve is established during embryogenesis, during which germ cells are limited to either entering meiosis, generating oocytes that remain arrested until puberty, or undergoing apoptosis (4). This limited number of oocytes shows signs of developmental decline with age in both mice and humans (5). Other animals such as the fruit fly instead produce oocytes throughout their lifespans due to a well-defined germ line stem cell niche that persists through adulthood (6). However, the mechanisms controlling oogenesis and the establishment of reproductive potential in other metazoans remain less understood. Moreover, numerous evo-devo studies have identified conserved genes expressed in germ line cells (7), but much less is known about the evolutionary history of the other somatic cell types within ovaries.

Echinoderms are non-chordate deuterostomes (8), a phylogenetic position informative for both vertebrates and other invertebrates, thus making them a valuable experimental system for addressing questions about the deep evolution of reproductive cell types (**Fig. 1A**). Echinoderms including sea stars continuously produce millions of oocytes that develop into optically transparent embryos and larvae, and have documented lifespans that can exceed a century (9). However, while echinoderms have been extensively used to study fertilization, gene regulation, and embryogenesis due to the abundance of eggs they provide, we still lack a mechanistic understanding for how they achieve this remarkable reproductive output (10,11).

**Figure 1.**
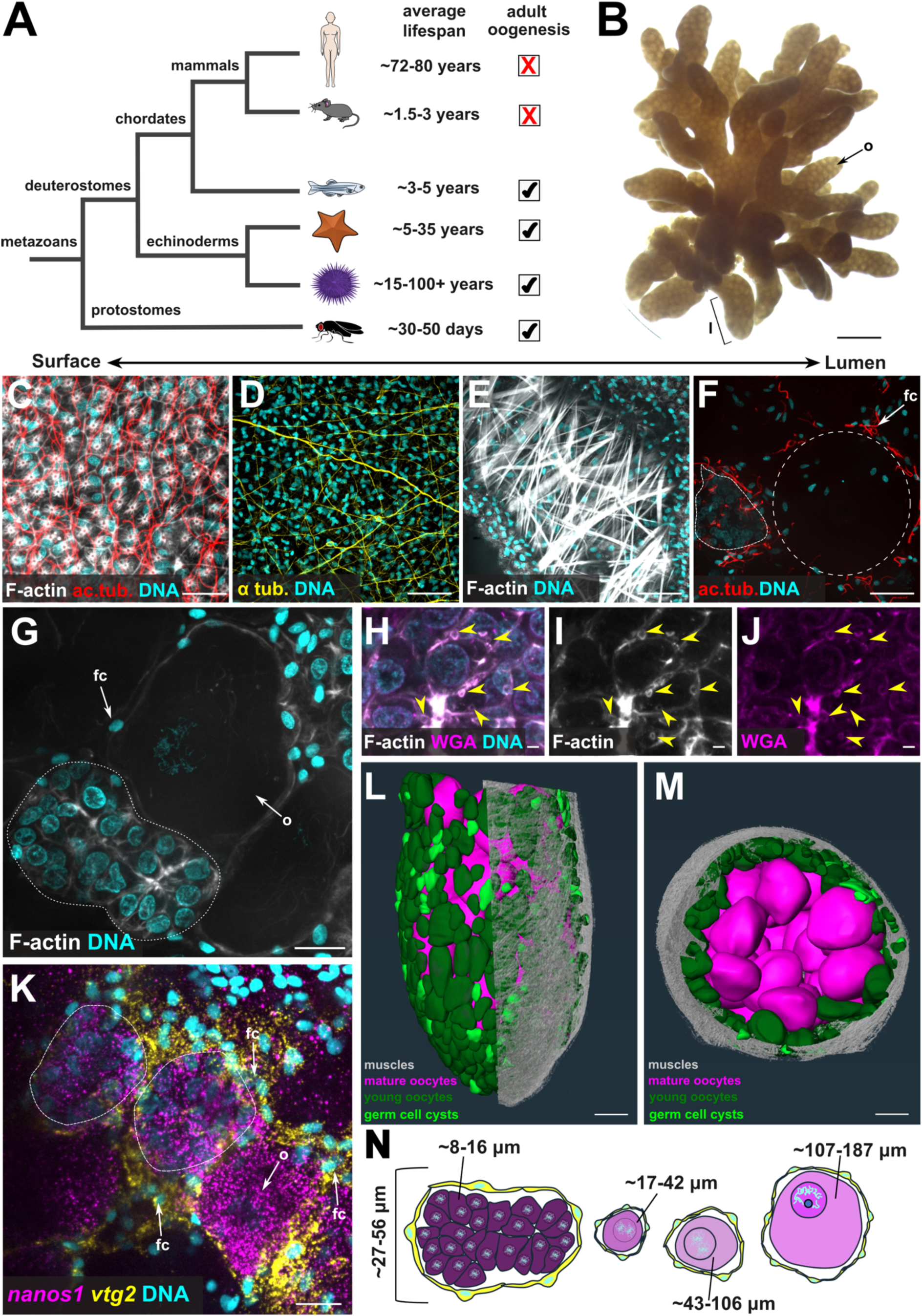
Molecular characterization of the *P. miniata* ovary. (**A**) Phylogenetic tree showing the evolutionary relationships of *P. miniata* with other metazoan experimental systems. (**B**) *P. miniata* ovarian explant. Scale bar is 1 mm. (**C**) Phalloidin and acetylated tubulin staining to label cell boundaries and cilia. Scale bar is 20 μm. (**D**) IHC for α-tubulin labeling the ovarian nerve plexus. Scale bar is 40 μm. (**E**) Phalloidin staining showing ovarian muscle fibers. Scale bar is 40 μm.(**F**) Acetylated tubulin labeling cilia on the follicle cells. Dotted circle indicates oocytes, while germ cell cysts are outlined with dotted lines. Scale bar is 20 μm. (**G**) Phalloidin staining labeling the cell boundaries of the germ cell cysts. Germ cell cysts are outlined with dotted lines. Scale bar is 20 μm. (**H-J**) Phalloidin and WGA lectin co-staining labeling the cell boundaries and membranes. Yellow arrowheads indicate intercellular bridge-like structures. Scale bar is 1 μm. (**K**) HCR for *nanos1* (magenta) and *vtg2* (yellow) labeling the entire germ lineage and follicle cells respectively. Germ cell cysts are outlined with dotted lines. Scale bar is 20 μm. Nuclei were labelled with DAPI (cyan). (**L-M**) 3D rendering of segmented germ cell cysts and oocytes. Scale bar is 100 μm. Ovary seen from the lateral (I) and top (J) views. (**N**) Schematic summary of the germ cell cyst, young oocyte and mature oocyte size distributions in the *P. miniata* ovary. fc, follicle cells; l, lobe; o, oocytes.

In this work, we asked how the reproductive potential of the sea star *Patiria miniata* is controlled, and whether sea star ovarian cell types are conserved with distantly related animals. Using high-resolution microscopy and single cell RNA sequencing (scRNA-seq), we defined the cellular organization of the sea star ovary, determined its cell types, and explored their evolutionary relationships with other animals. We found the ovary to consist of epithelial cells, immune cells, neurons, oocytes, three diversified groups of follicle cells, and putative oogonial stem cells organized in germ cell cysts. Our cross-species comparison revealed the presence of potentially conserved ovarian cell types, including a granulosa-like population. Last, we report conserved signaling pathways involved in oogenesis, as well as an intrinsic hypothalamic-like and gonadotropic-like signaling system operating between oogonial stem cells and ovarian neurons which we propose may be analogous, or possibly homologous, to the vertebrate HPG axis.

## Materials and Methods

### Animal husbandry, ovary isolation, *in vitro* culture

Adult *Patiria miniata* were obtained from South Coast Bio (San Diego, CA, USA) and Monterey Abalone Company (Monterey, CA, USA). The animals were maintained in temperature controlled (15°C) circulating seawater aquariums at the Marine Resources Center (MBL, MA, USA). Ovarian fragments were surgically isolated through the oral side of the sea star’s arm using a Stoelting™ Sterile Disposable Scalpel, No.11 and ovaries were collected using forceps in Eppendorf tubes containing filter sea water (FSW). For the single cell RNA sequencing experiments the ovaries were used immediately. For the rest of the experiments of the manuscript, ovaries were kept in FSW 10 μg/ml trimethoprim and 50 μg/ml sulfamethoxazole at 15°C as previously described (12).

### Ovary dissociation

Dissociation of isolated sea star intact ovarian lobes into single cells was performed as previously described with minor modifications (13). Briefly, ovarian fragments were transferred with forceps to a petri dish containing Ca^2+^ Mg^2+^-free artificial sea water (CaMg-free ASW) and allowed to settle for 3 minutes (min). Then the ovary was transferred to an Eppendorf tube containing the dissociation buffer (1M glycine, 0.02M EDTA, in CaMg-free ASW) and incubated for 10 min. Dissociation was promoted through gentle pipette aspiration every 2 min. Once dissociation was complete, cells were spun down with a swing bucket centrifuge at 500g for 10 min and washed twice with CaMg-free ASW. Propidium Iodide and Fluorescein diacetate were used to assess cell viability and only cell suspensions with cell viability ≥ 90 % were further processed. Single cells were passed through a 40 μm cell strainer to remove aggregates. The number of cells was estimated using a hemocytometer and diluted according to the manufacturer’s protocol (10X Genomics). During the entire process, specimens were kept on ice, except for the centrifugations in which they were kept at 4°C. The isolated cells were loaded on the 10x Genomics Chromium Controller according to the manufacturer’s instructions and cDNA libraries were generated using the Chromium Single Cell 3’ Reagent Kit (v3.1 Chemistry Dual Index). Specimens were sequenced by the Azenta Life Sciences (Burlington, MA) sequencing service using the Illumina NextSeq 550 with a maximum read length 2 × 150 bp at a resolution of 350M PE reads.

### Single cell RNA sequencing mapping of reads and data analysis

Single cells from ovaries originating from three different animals were processed using the 10X Genomics Chromium scRNA-seq capturing system. The libraries generated were biased towards early oogenesis since the 10X Genomics Chromium capturing system allows for the capture of cells up to 40 μm, which means that in our single cell atlas the maximum oogonium size represented is 40 μm. Fully grown oocytes in *Patiria miniata* are ∼170 μm. *Patiria miniata* genome version 3.0 (14) was used for single cell RNA sequencing read mapping. The corresponding annotation was modified to contain only protein coding entries using AGAT v0.7.0 (15) and Linux command line tools GNU coreutils v8.26. Cell Ranger Software Suite v7.1.0 (16) was used to create the index from the genome FASTA file and the annotation GTF file as well as for mapping with --no-bam and --nosecondary flags, with all other settings left as default so the number of cells was estimated automatically by Cell Ranger. Cell Ranger output files were loaded in R Studio and processed with the Seurat pipeline (17). Seurat objects were created by excluding cells containing genes that are transcribed in less than three cells and cells that have less than a minimum of 400 and a maximum of 5000 transcribed genes. Next the Seurat objects were normalized, and variable genes were found through the variance stabilizing transfer (VST) method with a maximum of 2000 variable features. The three different objects were integrated through identification of gene anchors (FindIntegrationAnchors). Following integration, scaling and principal component analysis (PCA) were performed. Jackstraw was used to evaluate PCA significance. Uniform Manifold Approximate and Projection (UMAP) was used to perform clustering dimensionality reduction. The final object consisted of 35,877 cells. The FindAllMarkers command was used to identify differentially expressed marker genes. The average expression of genes of interest were visualized using the DotPlot and DoHeatmap commands included in the Seurat R package. To compare the *Patiria miniata* ovary integrated single cell atlas with the publicly available human (18), mouse (19), zebrafish (20), sea urchin (21) and fruit fly (22) ovarian atlases as well as with the mouse pituitary gland atlas (23) and the sea urchin larva one (13) SAMap v1.02 (24) was used as previously described (25). For the mouse, zebrafish, fruit fly and sea urchin (ovary and larva) datasets the already available rds files were used, while for the human fetal ovary and the mouse pituitary gland datasets clustering analysis was performed similar to what was previously described (18,23). The average score of genes of interest was performed using the AddModuleScore function incorporated in the Seurat R package. The oogenesis trajectory was performed using the R package Monocle3 (26). The package EnhancedVolcano was used to visualize the differentially expressed genes between the germ cell cysts and oocytes along the trajectory, while the R packages topGO and clusterProfiler were used for the GO enrichment analysis.

### Light sheet microscopy/Segmentation 3D analysis

Ovarian samples were mounted individually in Fluidflon FEP tubes (Proliquid 1.6×2.4mm) using 1% low gelling agarose (Sigma-Aldrich A0701, type VII-A) with their elongated morphology parallel to the tube. A Zeiss Lightsheet 7 was used to acquire multiple-view data. It was equipped with a 10x/0.5 W Plan-Apochromat detection objective and two 10x/0.2 air illumination objectives producing two light-sheets 5.8 µm thick at the waist and 12µm thick at the edges of a 1100 µm x 1100 µm field of view. Illumination mode was dual (mean fused) with active pivot scan. Samples were imaged from five views (72 degrees apart) and Zoom factor 0.8. Each z-stack has a pixel frame of 1920×1920 and a voxel size of 0.57 µm x 0.57 µm x 1.8 µm. Each optical slice was acquired with a combination of 405, 488, and 561 nm laser with following filter settings: 405nm: EF1 - BP 420-470, CAM BS - SBS LP 490; 488 nm: EF1 - BP 505-545, CAM BS - SBS LP 560, 561 nm: EF1 - BP 575-615, CAM BS - SBS LP 640. For registration of multi-view acquisitions and fusion plus deconvolution of the registered data, the Fiji plugin BigStitcher v2.3.4 (27) was used with standard settings. Image segmentation, 3D-reconstructions and data analysis were performed using the Amira software package (FEI Visualization Sciences Group Thermo Fisher Scientific), version 2024.1. Morphological features such as muscles and nuclei were segmented semi-automatically using various tools in Amira, such as “Thresholding” and “Magic Wand”. Oocytes and germ cell cysts were primarily segmented manually. Volumes of oocytes and germ cell cysts were measured using the “Material Statistic” tool. Visualization of all segmented reconstructions was achieved using tools like “Volren”, “Surface Rendering” and “Surface View” within the software package.

### Immunohistochemistry (IHC) and fluorescent stainings

Ovarian fragments were fixed with 4% PFA in FSW for 15-20 min at room temperature (RT). Specimens were washed twice with FSW and five times with PBS supplemented with 0.1 % Tween (1x PBST). Next, samples were incubated in blocking solution [1 mg/ml Bovine Serum Albumin (BSA) and 4 % sheep serum (SS)] in PBST for one hour at RT. Blocking solution was removed and primary antibodies were added in the appropriate dilution in blocking solution and incubated overnight at 4°C. 1E11 (gift from Dr. Ina Arnone) was used in a 1:20 dilution, anti-acetylated tubulin (Sigma) 1:100, anti-DM1α (Developmental Studies Hybridoma bank) 1:100, anti-alpha tubulin (Sigma) 1:100, anti-MHC (PRIMM) 1:50, anti-serotonin (Sigma) 1:500, anti-TH (Sigma) 1:100m and anti-Myosin II (Developmental Studies Hybridoma bank) (1:50). Samples were washed five times with 1x PBST (15 min each) and incubated for one hour at RT with secondary antibodies (AlexaFluor) diluted 1:1000 in PBST. Samples were washed five times with 1x PBST. In cases where phalloidin (Invitrogen) was used, it was diluted 1:20 (from a 40x stock solution) in 1x PBST and allowed to incubate overnight at 4°C. For WGA staining (Invitrogen) a working solution of 5 μg/mL was prepared, and the specimens were incubated overnight at 4°C. Ovaries were washed five times with 1x PBST, nuclei were stained with DAPI (1 μg/mL), and specimens were mounted for imaging. During the entire procedure the interval of each wash was 15 min.

### Whole mount Hybridization Chain Reaction (HCR)

HCR was performed according to previous protocols (28) with minor modifications. Probes were designed either by Molecular Instruments or HCR 3.0 Probe Maker (29). Ovarian fragments were fixed in 4% PFA in Fixative Buffer (0.1M MOPS, 0.5M NaCl, 2mM EGTA, 1mM MgCl_2_ and 1x PBS) for 1 hour at room temperature (RT). Specimens were washed three times with Fixative Buffer (without PFA) at RT. Samples were washed twice with 100% methanol, transferred into fresh 100% methanol, and placed at −20°C for long term storage. Specimens were rehydrated in 75%, 50%, 25% methanol in RNAase free water on ice (each for 10 min). Next, ovaries were washed four times for 5 min each in RNase free 1x PBST. Ovaries were treated with permeabilization buffer (1.0% SDS, 0.5% Tween, 50 mM Tris-HCl (pH 7.5), 1.0mM EDTA (pH 8.0), 150 mM NaCl) and incubated for 1h at RT. Next, specimens were washed four times for 5 min each in 1x PBST. Pre-heated (37°C) Probe Hybridization Buffer (PHB) was added to the samples and incubated for 3 hours at 37°C. PHB was exchanged with PHB containing in total 0.04 μM of each probe and specimens were left to incubate for one day at 37°C. Following hybridization, samples were washed 4 times with pre-heated Probe Wash Buffer (PWB) for 5 min each, followed by two washes with PWB for 30 min each at 37°C. Samples were then washed two times with 5x SSCT (5x Saline Sodium Citrate and 0.1% Tween) and two times with 1x SSCT (1x Saline Sodium Citrate and 0.1% Tween) each for 5 min. Next, specimens were incubated with Amplification Buffer for 30 min at RT, hairpins were added to the specimens according to manufacturer’s guidelines and incubated overnight at RT in the dark. Lastly, samples were washed twice with 1x SSCT for 5 min, two more times for 30 min and four times with 1x PBST. Nuclei were stained with DAPI (1 μg/mL), and specimens were mounted for imaging.

### Fluorescent *in situ* hybridization (FISH)

FISH and antisense RNA probes design was performed as previously described (30,31). In brief, ovaries were fixed in 4% PFA in MOPS Buffer for 1 hour at RT. Samples were washed five times with MOPS buffer, gradually dehydrated (20%, 50% ethanol) and stored in 70% ethanol and kept at −20°C for long term storage. Specimens were gradually rehydrated (50%, 20% ethanol) and washed five times with MOPS Buffer. At this step ovaries were dissected to smaller pieces using micro scissors. Specimens were then pre-hybridized for 3 hours at 65°C, followed by a 48 hour period of probe hybridization. Next, specimens were washed five times with MOPS Buffer and incubated in blocking solution (Akoya Biosciences) for 30 min at RT. Next, specimens were transferred to a solution containing the anti-Digoxigenin (Roche) antibody diluted 1:1000 in blocking solution. The antibody solution was removed, samples were washed with MOPS Buffer five times and the signal was developed using fluorophore conjugated tyramide technology (TOCRIS). Nuclei were stained with DAPI (1 μg/mL), and specimens were mounted for imaging.

### EdU labeling

Click-It EdU Cell Proliferation Kit for Imaging Alexa Fluor 488 (Thermo Fisher Scientific) was used and was combined with either HCR or IHC as previously described (13,32). Isolated intact ovarian fragments were treated with EdU at a final concentration of 20 μM in FSW with 10 μg/ml trimethoprim and 50 μg/ml sulfamethoxazole and allowed to incubate for 2 days at 15°C. Depending on whether the EdU cell proliferation was combined with HCR or IHC specimens were fixed accordingly as described in their respective sections above. To develop the EdU signal, the Click-iT reaction mix was prepared according to the manufacturer’s guidelines. In cases where the EdU assay was combined with HCR after the washes with PWB, specimens were washed twice with 3% BSA in PBST for 15 min at RT. Next the Click-iT reaction mix was added and incubated for 30 min in the dark at RT. After this step the HCR procedure was carried out as described above. For combination of the EdU Cell Proliferation assay with immunostaining the ovaries were fixed in 4 % PFA in FSW for 15 min at RT and washed five times with PBST. PBST was removed and the reaction mix was added to the samples for 30 min (RT). Specimens were washed five times with PBST, mounted for imaging.

## Results

### The structural organization of the *Patiria miniata* adult ovary

We first sought to define the structure of its ovary, which is multi-lobed and optically translucent (**Fig. 1B**). We stained for F-actin with phalloidin as a cell boundary marker and acetylated tubulin to visualize cilia (**Fig. 1C**). Each ovarian lobe is a hollow tube that consists of a thin, tightly packed epithelial cell layer with long motile cilia (**Fig. 1B**). We also identified cells organized in plexus-like structures resembling a nerve net underneath the epithelial layer that are positive for synaptotagmin 1, a pan-neuronal marker in echinoderms (**Fig. 1D**, **S1A, E-H**). We also identified ovarian neurons in the related sea urchin *Strongylocentrotus purpuratus* (**Fig. S1I-J**) and the sea cucumber *Holothuria tubulosa* (**Fig. S1K-L**), arguing that these are a conserved cell type across echinoderms. We also identified extensive musculature below and in close proximity to the nerve net (**Fig. 1E, S1B**), which is likely responsible for the contraction of the ovary during spawning (33). We observed axonal projections in close proximity to the muscle layer, which may regulate their contraction (**Fig. S1E-F, I-J**). Below the muscle layer and within the lumen, we detected oocytes at different stages of maturation enwrapped by follicle cells, which are filamentous in nature and ciliated (**Fig. 1F, Fig. S1C-D**).

We determined the stages of oocyte development, and the nature of oocyte-follicle complexes. Early small oogonia are spatially organized in closely associated groups with an average size of 8-16 μm (**Fig. 1G, Fig. S2A-B**). These groups were found in all ovaries tested, although their number varied with different individuals. The chromatin morphology of these cells was distinct, with condensed DNA concentrated towards the nuclear periphery. This may be consistent with the leptotene to zygotene stage of meiosis, in which chromosomes adopt a bouquet configuration (34). The cells within these groups were compact with intense actin staining at the cell boundaries, and surrounded by small somatic nuclei that do not intercalate within the cluster (**Fig. 1G**). This organization resembles germ cell nests or cysts that are found in other species, including fruit flies and mammalian fetal ovaries (35). Similar structures were observed by electron microscopy in the ovaries of the phylogenetically distant sea star *Asterias rubens* (36). Co-staining for F-actin and membrane lectins revealed cortical channel-like structures between distinct oogonia within the germ cell groups (**Fig. 1H-J**). Such an organization strongly resembles the intercellular bridges that physiologically connect the germ cells within the germ cell cysts in mammals and protostomes, allowing for the exchange of the germ plasm and suggests that germ cells are organized in cysts also in a non-chordate deuterostome. Therefore, from here on we refer to these groups as “cysts”.

To test the germline nature of these cysts, we performed HCR for the conserved germ cell factor *nanos1,* and the yolk protein *vtg2*, which is expressed in sea star follicle cells in other echinoderms (37–39). We found that *nanos1* is expressed specifically in oocytes of all sizes as well as in the cysts (**Fig. 1K**). *vtg2* was expressed by the follicle cells that surround the oocytes. We also found that the *vtg2+* follicle cells enwrap the cysts as a whole, while none of the cells within the cysts are individually enwrapped (**Fig. 1K**). This configuration further resembles the germ cell cysts observed in vertebrates, suggesting that sea star adult oogenesis may share conserved features. Co-staining for *nanos1* and *vtg2* also enabled us to ask what is the smallest oocyte size necessary to form a primary follicle. We found that the smallest *nanos1+* oocytes are located next to germ cell cysts and by using the diameter of these oocytes as a proxy to calculate the size and volume we found they are within the range of 17-42 μm (**Fig. S2C-D**).

We next asked how oogenesis is spatially organized within the ovarian lobe. Is there a defined stem cell niche, or is oogenesis instead uniformly distributed? To test this, we used large volume light sheet microscopy on entire ovarian lobes (**Fig. 1L-M, Fig. S2G**). Segmentation and 3D reconstruction revealed that oogenesis is spatially polarized from outside to inside. Germ cell cysts and smaller oocytes were localized entirely to the periphery of the lobe interior, but were broadly distributed. Fully grown oocytes in prophase I arrest, instead, were centrally located within the lumen (**Fig. 1L-M, Fig. S2G**). Quantification of segmented cell volumes found that the diameter of the entire germ cell cysts ranges from 27-56 μm, of the small oocytes from 17-106 μm and of the pre-GVBD ones from 107-187 μm (**Fig. S2E-F**). Altogether, these results define the cell type organization of the sea star ovary at a molecular level and the oocyte size distribution during oogenesis (**Fig. 1N, Fig. S2A-F)**. Moreover, our data provide molecular evidence for the presence of germ cell cysts and show how *P. miniata* oogenesis is spatially oriented within the ovary.

### The *P. miniata* ovary consists of nine molecularly distinct cell type groups

Having defined the overall structures of the ovary, we next sought to determine the molecular identities of its constituent cell types with scRNA-seq (**Fig. 2A, Fig. S3A**). We used ovaries from three independent *Patiria miniata* females (**Fig. S3B-C**). One ovary originated from an individual in which spawning was induced 2 months prior to ovarian isolation, one was highly enriched in germ cell cysts and early oocytes, and one consisted mostly of fully grown oocytes. For each ovary used, the size distribution of their oogonia/oocytes was determined by dissociating the ovaries and manually measuring the oogonia/oocytes diameter to estimate their volume (**Fig. S3B-C**). Computational analysis and integration of the sequenced libraries yielded an atlas of 35,877 cells distributed in twenty-seven clusters or cell states (**Fig. 2A, Fig. S3A**).

**Figure 2.**
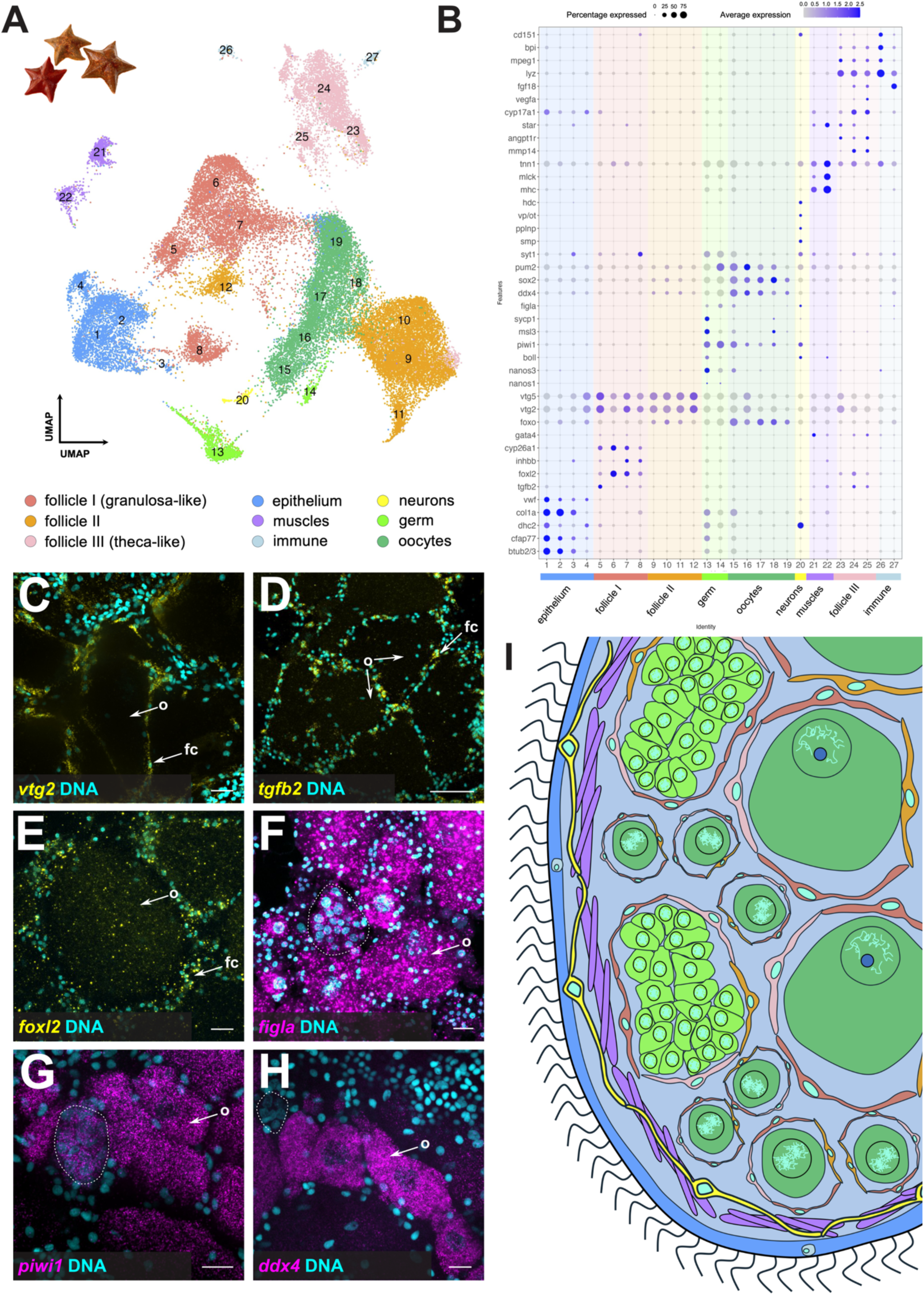
Cell type/state diversity of the *P. miniata* adult ovary. **(A)** Integrated UMAP of the three different scRNA-seq libraries. Color code is indicative of different cell type groups. **(B)** Dotplot showing the average expression of known echinoderm cell type markers as well as genes labeling ovarian and associated cell types in other organisms. Color code is the same as in panel A. **(C-D)** HCRs for the follicle cells markers *vtg2* (C) and *tgfb2* (D). **(E-F)** FISH using antisense probes for the transcription factors *foxl2 (E)* and *figla* (F) labelling the follicle cells and germ lineage respectively. Dotted circle in panel E is indicative of oocytes, while germ cell cysts in panel F are outlined with dotted lines. Scale bars for panels C and E are 20 μm, and for panel D 60 μm. **(G-H)** HCRs for the germ line markers *piwi1* and *ddx4*. Germ cell cysts in panels G and H are outlined with dotted lines. Scale bars are 20 μm. **(I)** Schematic representation of the ovary depicting the cell type groups revealed by our single cell RNA sequencing atlas. Color code is the same as in panel A. fc, follicle cells; o, oocytes.

We next inferred the cell type identities of the twenty-seven clusters using the average expression of differentially expressed marker genes, and gene homologs known to label ovarian cell types, in echinoderms and other organisms. (**Fig. 2B)** We curated 9 cell type groups expressing genes corresponding to endothelial and ciliated cells (*btub2/3, cfap77, dhc2, col1a, vwf*) (13,40,41), mammalian granulosa cells (*tgfb2, foxl2, inhbb, cyp26, foxo, gata4*) (1,42–45), echinoderm follicle cells (*vtg2, vtg5*) (39), mammalian theca cells (*mmp14, angpt1, star, cyp17a1, vegfa, fgf18*) (46–50), muscles (*mhc, mlck, tnn1*) (13), immunocytes (*lyz, mpeg1, bpi, cd151*) (51–54), neurons (*syt1, smp, pplnp, oxy/vp, hdc*) (13,55,56) and germ cells/oocytes (*nanos1, nanos3, boll, piwi1, msl3, sycp1, figla, ddx4, sox2, pum2*) (37,38,57–64). We found that most cell states were equally distributed amongst the three ovary samples (**Fig. S3D**) with the exception of clusters 5, 9, 10, 11, 12 and cluster 4, identified as follicle cells and epithelium, respectively, which predominantly originated from the previously spawned ovaries of female 1. Germ cell cyst clusters 13 and 14 were also enriched in sample 2, which was used for the 3D volume analysis (**Fig. 1I-K**) and contained numerous germ cell cysts. Therefore, our single cell atlas reflects the experimentally validated reproductive states of the ovaries used. With this confidence in our dataset, we next sought to spatially map their location within intact ovaries using HCR and FISH (**Fig. 2C-H. Fig. S3E-I**).

We first examined genes expressed in follicle cells, which enwrap and support oocytes as they develop. Using HCR, we detected *vtg2* (a previously validated marker) and the signaling molecule *tgfb2* in follicle cells, with the mammalian theca marker *tgfb2* (65) being expressed in a subset of them (**Fig. 2C-D**). We also detected *foxl2*, a known mammalian granulosa cell transcription factor, in follicle cells (**Fig. 2E**). Therefore, the clusters that contain *foxl2* (clusters 5-8, 23-24), *tgfb2* (clusters 5, 7-8, 22, 24-25) and *vtg2* (clusters 5-8, 9-12, 23) transcripts were confirmed to correspond to the partially non overlapping follicle cell groups as identified by scRNA-seq.

We next turned to the putative germ line clusters. The transcription factor *figla*, which functions in mammalian oogenesis, was detected exclusively in the germ cell lineage including cysts and oocytes (**Fig. 2F**). Similar results were obtained for the germline factors *piwi1* and *nanos3* (**Fig. 2G, Fig. S3D-D’**). Surprisingly, we did not detect the conserved germ line RNA helicase *ddx4/vasa* in the germ cell cysts, though it was detected in all other oocyte stages (**Fig. 2H**). This result suggests that *ddx4/vasa* transcription is activated only once the primary follicle is formed. These expression patterns are consistent with the single-cell predictions and suggest the presence of two cell groups: the germ cell cysts (clusters 13 and 14) and early oocytes (clusters 15, 16, 17, 18 and 19). To further validate our clustering analysis, we examined *syt1* expression, which was unexpectedly predicted to be present not only in the nervous system but also in germ cell cysts and oocytes. HCR confirmed *syt1* in the nerve net as well as in the germ cell cysts and oocytes (**Fig. S3F-G’**), while a positive control in *P. miniata* larvae and post-metamorphic juveniles verified the probe specificity, showing *syt1* expression in the nervous system as reported in other echinoderms (**Fig. S3H, I**). Taken together, our spatial and transcriptomic analyses enabled us to identify and map nine cell type groups distributed in twenty-seven clusters (**Fig. 2I**).

### Shared cell type toolkits between *P. miniata* and other deuterostomes

How similar are sea star ovarian cell types to those of animals that diverged millions of years ago? Our gene candidate-based analysis identified similarities—for instance, sea star follicle cells (clusters follicle I and follicle III) express gene orthologs also found in mammalian granulosa (*foxl2, inhbb, tgfb2, gata4*) and theca (*tgfb2, mmp14, angpt1r, star, cyp17a1, vegfa, fgf18*) cells. To more broadly test this similarity, we used SAMap to compare our atlas with published datasets of adult fruit fly, sea urchin, zebrafish, mouse, and fetal human ovaries (**Fig. 3, S4**). As expected, sea star cluster 22 (muscle-related) aligned with muscle clusters in all animal ovaries tested. This similarity is not unique to ovarian muscles, as alignment with a sea urchin larval scRNA-seq dataset also revealed strong similarity with esophageal muscles (**Fig. 3, S4**). We also identified similarity between the sea star follicle cell I (cluster 6) and their follicular counterparts in fruit fly and sea urchin (**Fig. 3A, S4A**). Surprisingly, despite substantial evolutionary distance, this follicle cell cluster also aligned with follicle cells in zebrafish, and granulosa cells in both mouse and human, although in the latter with lower alignment scores (**Fig. 3B,C, S4B**). In addition, the sea star oocyte cluster 19 aligns with germ and oogonial cell clusters in fruit fly, sea urchin, zebrafish and human (**Fig. 3A-C, S4A**). Zebrafish and mouse theca cells aligned with sea star epithelial clusters, while zebrafish vasculature aligned to one sea star follicle cluster (**Fig. 3B**). Similarities become scarcer when we compare the ovaries of sea star with the adult mouse, which does not contain oogonial stem cells (66) (**Fig. S4B**). However, when we compared the sea star ovary with the human embryonic ovary, in which oogenesis is still occurring, we found that one human germ cell cluster aligned with the sea star oocytes cluster 17 (**Fig. 3C**). We also detected similarities between sea star follicle cells (I) with human endothelium, sea star follicle cells (III) with human granulosa cells, as well as sea star epithelium with human stromal and theca cells (**Fig. 3C**). A summary of the main findings can be found in (**Fig. 3D**). In support of the specificity of the similarities we detected, comparison to the sea urchin larva, which serves as a null comparison, yielded no strong hits apart from muscle (**Fig. S4C**). Overall, our analysis has shown the existence of transcriptionally similar ovarian cell type programs across phylogenetically distant organisms, suggesting the possible pre-chordate origins of ovarian cell types, including follicle cells.

**Figure 3.**
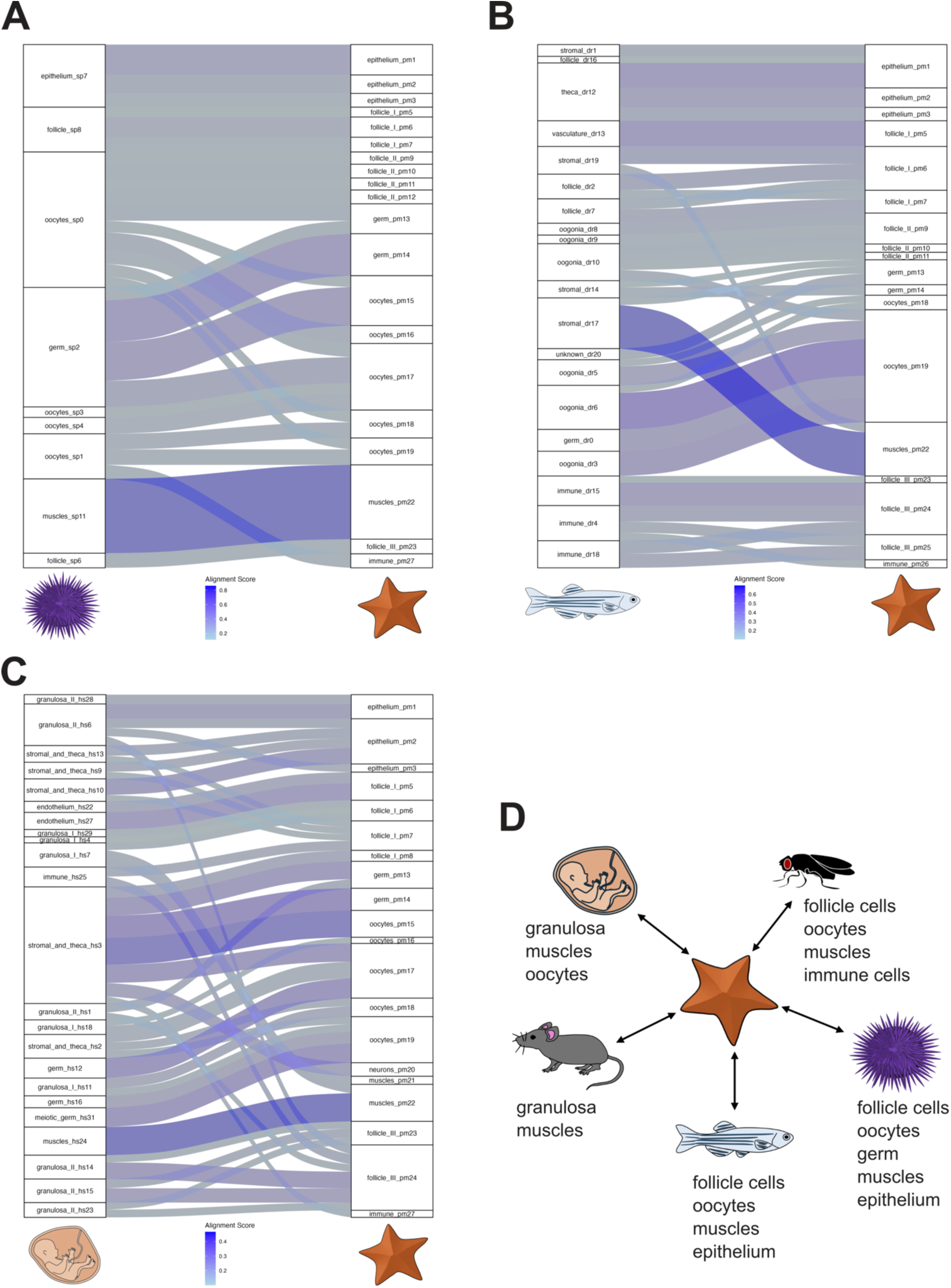
Cross-species comparison of ovarian single cell atlases with SAMap. Sankey plot depicting the alignment scores between the adult sea star ovarian cell clusters with the adult sea urchin **(A)**, adult zebrafish **(B)** and fetal human ovarian clusters **(C)**. **(D)** Summary of the cell types shared between sea star and the taxa used for comparison. Alignment score is defined as the average number of mutual nearest cross-species neighbors of each cell relative to the maximum possible number of neighbors. Alignment scores equal or greater to 0.1 were considered significant.

### Identification and characterization of mitotic oogonial stem cells

Having defined similar ovarian elements across species, we then asked what unique features of the sea star ovary support its extreme reproductive capability, such as adult oogonial stem cells (OSCs). To identify potential OSC populations, we computationally scored for the co-expression of cell cycle-associated and meiotic genes with germline marker genes (**Fig. 4A, S5A**). We found that the epidermal cell clusters 2 and 3, the germ cell cysts clusters 13-14 and oocytes (15–19) express genes associated with G1/S phase (including *cyclinD2, cyclinE, cdt1* and *pcna*), while the epidermal cluster 4, the follicle cell clusters 9-12, the germ cell cyst cluster 14 and oocytes (15–19) express genes operating during G2/M phase (including *cyclinA, cyclinB* and *bub3*) (**Fig. 4A**). Genes specific to meiosis, such as *spo11, mlh1, hormad, sycp1, sycp2* and *sycp3* were expressed in the germ cell cyst clusters 13-14 and oocyte clusters 15-19 (**Fig. 4A**). HCR for one of those genes, *mlh1,* confirmed the predicted expression in both the germ cell cysts and the oocytes (**Fig. 5SB**).

**Figure 4.**
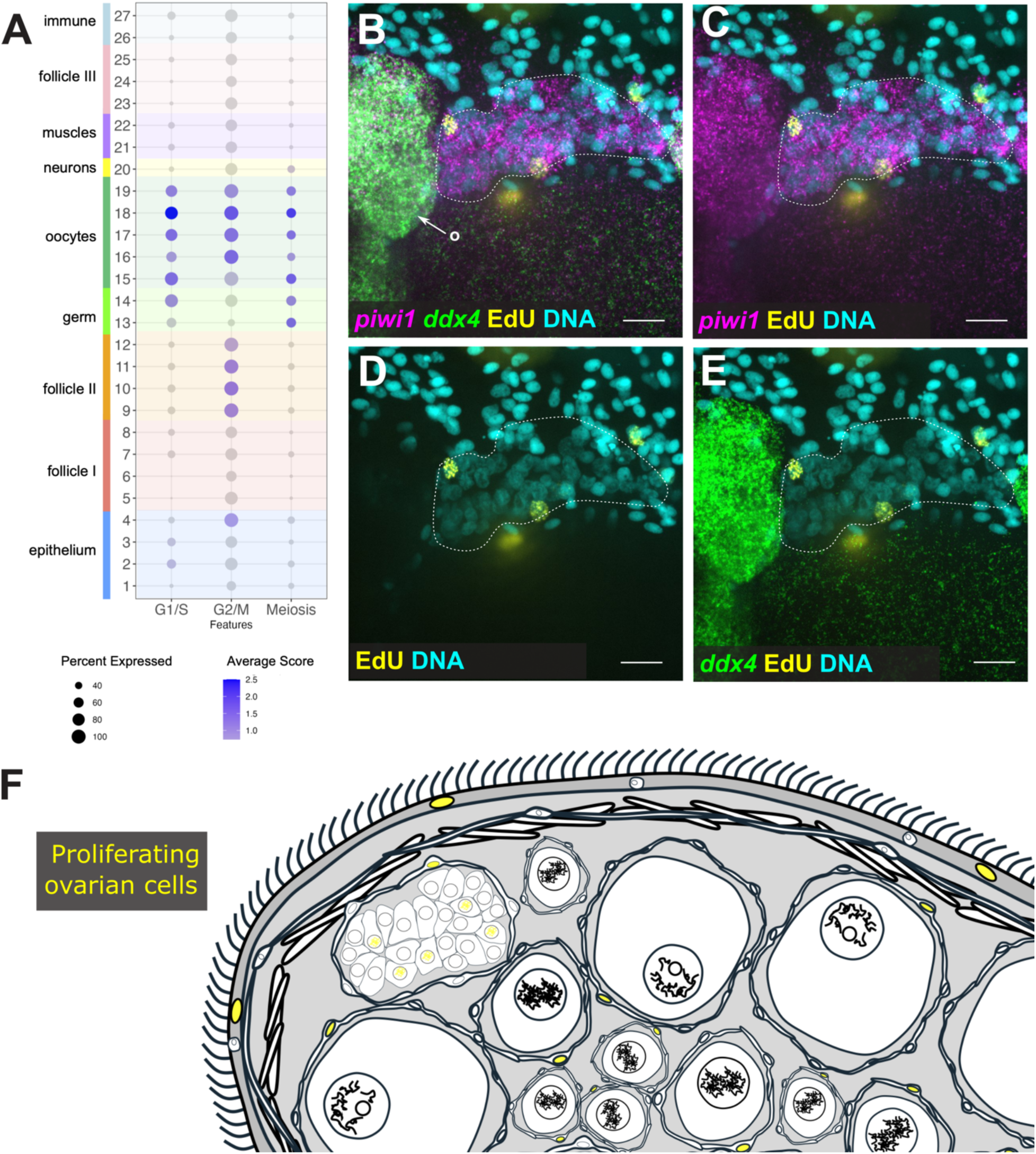
Identification of self-renewing oogonial stem cells. **(A)** Dotplot showing the average score of genes involved in G1/S, G2/M phases of the cell cycle and meiosis across the *P. miniata* ovarian cell type groups. (**B-E**) HCR for the germline markers *piwi1* and *ddx4* paired with the EdU proliferation assay. Scale bars are 20 μm. Nuclei were labelled with DAPI (cyan). Dotted lines outline the germ cell cyst in panels B-D. **(F)** Summary of the ovarian cell types undergoing mitosis as revealed by the EdU nuclear incorporation. o, oocytes.

**Figure 5.**
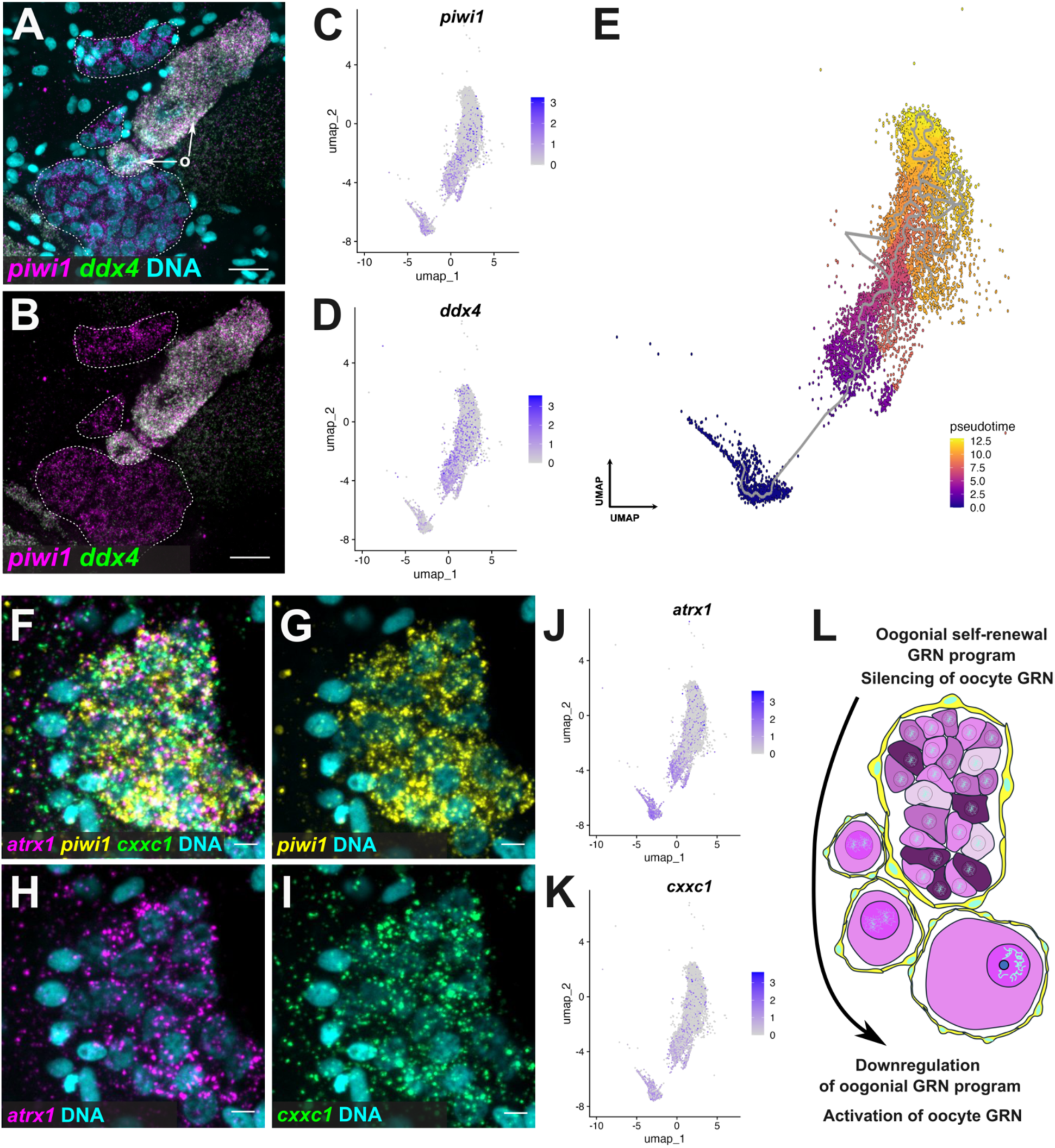
Reconstruction of the early oogenesis developmental trajectory. **(A-B)** HCR for the germline markers *piwi1* and *ddx4* highlights the absence of *ddx4* in the germ cell cysts. Dotted lines outline the germ cell cyst. Nuclei were labelled with DAPI (cyan). **(C-D)** Feature plot showing the expression levels of *piwi1* and *ddx4* along the germline clusters of the *P. miniata* single cell atlas. **(E)** UMAP depicting the developmental trajectory of the germline clusters reconstructed with Monocle3. The germ cell cluster 13 has been used as a root. **(F-I)** HCR for the transcriptional regulators *atrx1* and *cxxc1* paired with *piwi1* confirms their expression in germ cell cysts. Scale bars are 5 μm. **(J-K)** Feature plot showing the distribution of *atrx1* and *cxxc1* transcripts along the trajectory confirms their differential expression in clusters corresponding to germ cell cysts. Nuclei were labelled with DAPI (cyan). **(L)** Cartoon highlighting the difference in gene expression of the oogonia in the germ cell cysts versus the oocyte stages. o, oocytes.

To directly test cell proliferation, we cultured ovarian lobes *in vitro* and performed EdU cell proliferation assays (**Fig. 5SC-E**). We found cells undergoing DNA replication in both somatic and germ cell populations. This included the ovarian epithelium, follicle cells, and within germ cell cysts (**Fig. 5SC-E**). To further define the germ line population, we paired EdU incorporation with HCR for *nanos1, piwi1* and *ddx4/vasa* (**Fig. 4B-E**). The EdU signal was exclusively detected in *nanos1* and *piwi1* positive germ cell cyst nuclei, but not in the *ddx4* positive growing oocytes. Altogether, these data revealed the presence of replicating cells within the sea star ovary (**Fig. 4F**) and support our hypothesis of a self-renewing oogonial stem cell pool in adult sea star ovaries.

We next asked how gene expression changes as oocytes develop. We used Monocle3 to identify a developmental trajectory linking the 7 germ line cell clusters (13, 14, 15, 16, 17, 18), from OSC to oocyte. Cluster 13 was chosen as the root because its molecular signature and cellular morphology was most similar to typical germ cells, and because it did not express *ddx4*, as experimentally validated by HCR (**Fig. 5C-E**). We found a substantial number of genes that are upregulated in germ cell cysts and downregulated in oocytes, and fewer that are more highly expressed in oocytes. (3378 genes upregulated in cysts, 849 upregulated in small oocytes; **Fig. S6A**). Gene ontology (GO) enrichment analysis shows that most of the cyst genes correspond to transcription and translation factors, while many of the oocyte genes are related to splicing, protein synthesis and transport. (**Fig. S6B-B’, S7**). Among the genes also upregulated in the germ cell cyst clusters are the transcriptional and epigenetic regulators *atrx1* and *cxxc1*. Both genes are important for mammalian sexual differentiation and oogenesis respectively (67–69). Plotting their average expression on the trajectory as well as HCR for both genes confirmed their co-expression in the oocyte lineage and presence within the germ cell cysts (**Fig. 5F-K, Fig. S6C-C’’’**). Taken together our data highlight, for the first time in echinoderms, distinct transcriptional differences occurring during the transition from oogonia to early oocytes (**Fig. 5L**).

### Signaling pathways across ovarian cell types

What are the major signaling pathways that could control oogenesis and folliculogenesis? We first mined our atlas for the expression of gene groups corresponding to either ligands or downstream signaling cascades of the Wnt, Tgfb, Delta/Notch, Hh, and RA signaling pathways (**Fig. 6A, S8A**).We found that the clusters that most strongly express Wnt ligands correspond to epithelium, oocytes and immune cells, while follicle I, germ, oocytes, neurons and muscles are able to respond to the signal based on the expression of frizzled receptors, *dvl3, gsk3, apc, tcf/Lef* and *β-catenin* (**Fig. 6A, S8A**). Similarly, Tgfβ ligands are expressed by the epithelium, follicle I, follicle III and muscle clusters, while their receptors and signaling cascade factors are expressed in germ, oocytes, muscles, follicle III and immune cells (**Fig. 6A, S8A**). For Delta/Notch signaling, *delta* transcripts are found in epithelial, follicle II, follicle III and oocytes cell type groups, and the signal is predicted to be received by follicle I, oocytes and neurons *notch* receptors and gamma secretase subunits (**Fig. 6A, S8A**). Hedgehog (*hh*) is exclusively expressed by neurons, and the signal may be received by follicle I and germ cell type groups based on the expression of patched and smoothened receptors (**Fig. 6A, S8A**). Lastly, retinoic acid synthesis machinery gene orthologs, including *sdr7, aldh1, aldh3* and *aldh8* are differentially expressed in the follicle II, germ, and oocyte cell groups, and the cells that express RA receptors *rar, rxr* and *ppar* belong to the follicle I, III, germ, neural and muscle ones (**Fig. 6A, S8A**). Degradation of RA is performed by cells expressing *cyp26,* which includes the follicle I group, while its transport is mediated by cells that express *fabpl3,* which includes the follicle I and follicle II groups (**Fig. 6A, S8A**). Our results indicate the presence of a complex signaling system involved in sea star oogenesis by which diverse signals are produced and received both by the follicle cells, the cells of the germ lineage, and the ovarian neurons.

**Figure 6.**
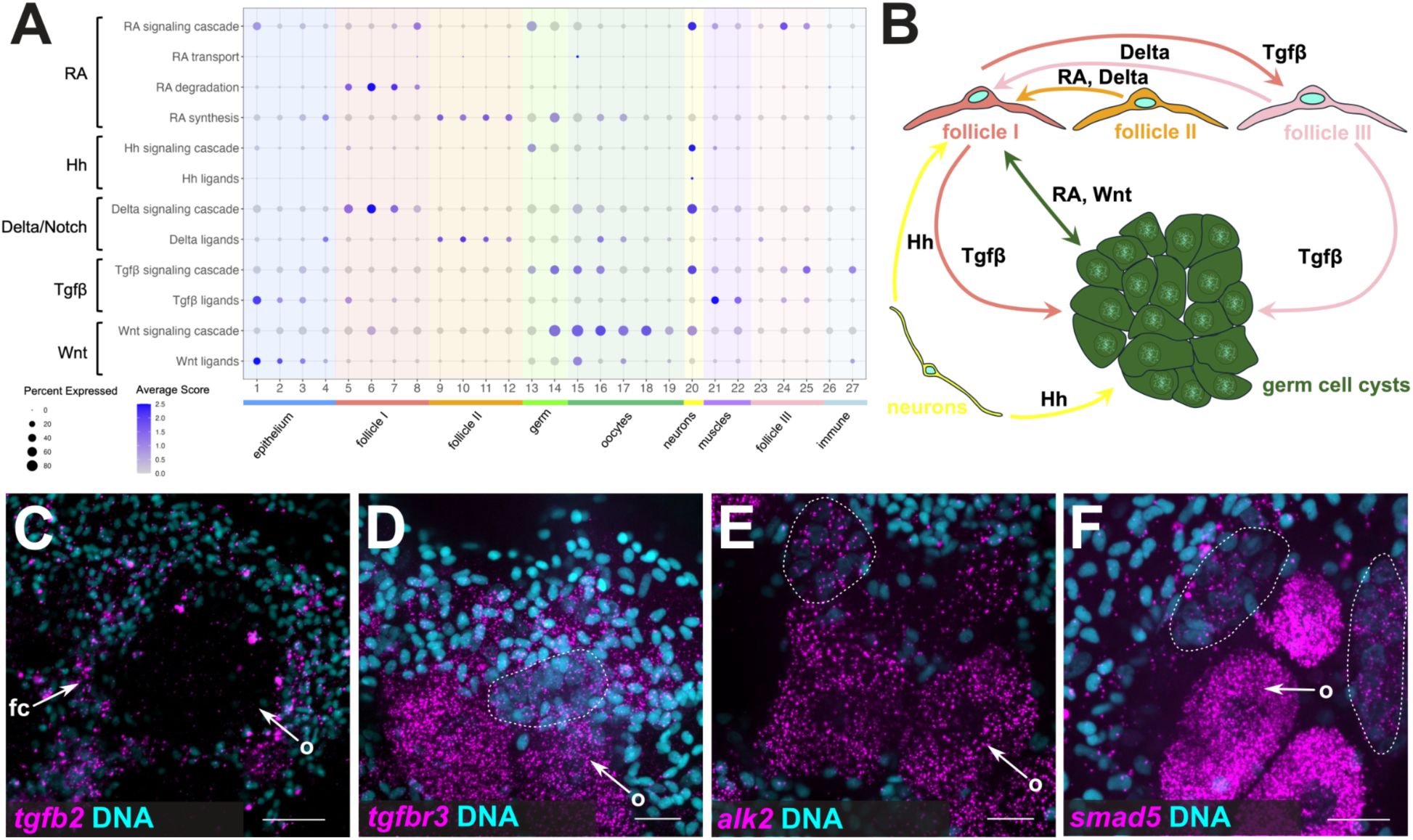
Putative signaling pathways involved in ovarian cell type crosstalk. **(A)** Dotplot showing the average score of gene groups corresponding to ligands and signaling cascade genes in relation to Wnt, Tgfβ, Delta/Notch, Hh and RA signaling pathways across the *P. miniata* ovarian cell type groups. (**B**) Schematic representation of the predicted paracrine crosstalk between follicle cells, neurons and germ cell cysts. (**C**) HCR for *tgfb2* labeling follicle cells. Scale bar is 40 μm. (**D-F**) HCRs for *tgfbr3*, *alk2* and *smad5* show localization within the germ cell cysts and oocytes spatially confirming the scRNA-seq predictions. Scale bar is 20 μm. Dotted lines outline the germ cell cysts. Nuclei were labelled with DAPI (cyan). fc, follicle cells; o, oocytes.

We next focused on Tgfβ signaling, which has been implicated in primordial germ cell development and follicular growth in mammals (1,70,71) (**Fig. 6C-F, S8B-D**). Our atlas indicates that Tgfβ signaling may be active within the oocyte-follicle complex. Of the genes tested, only *tgfb2* ligand was uniquely expressed in somatic tissues and primarily in follicle cells enwrapping the oogonia and mature oocytes (**Fig. 6C**). We found Tgfβ receptors *tgfbr3*, *alk2* and the intercellular messenger *smad5* expressed by both the germ cell cysts and developing oocytes, validating the scRNA-seq predictions (**Fig. 6D-F**). In addition, our atlas showed that several Bmp ligands are expressed within the germ lineage, suggesting that Tgfβ signaling may also be active between growing oocytes, or acting upon follicle cells, a dual role also reported in mammals (**Fig. 6A, S8A**). HCR for *bmp1* and *bmp4* confirmed these predicted expression patterns (**Fig. S8B-D**). In summary, our results support the deep evolutionary conservation of Tgfβ signaling pathway components within the ovarian follicle.

### Evidence for pre-chordate origins of the HPG axis

Vertebrate oogenesis is regulated by endocrine signaling between the hypothalamus, the pituitary gland, and the ovary. Therefore, we asked how the ovarian nervous system might serve as an intrinsic paracrine regulator of oogenesis. We found that most neuropeptide synthesis genes are specifically expressed by neuronal cluster 20, including, tryptophan hydroxylase (*tph*), tyrosine hydroxylase (*th*), choline O-acetyltransferase (*chat*), histidine decarboxylase (*hdc*), and glutamate decarboxylase (*gad*) are expressed, indicating serotonergic, dopaminergic, cholinergic, histaminergic and GABAergic neurons, respectively (**Fig. S9A**) (72,73). Moreover, we found that sixteen different neuropeptides are expressed in the same cell cluster (**Fig. S9A**). Immunofluorescence for TH and serotonin (5HT) confirmed the presence of extensive dopaminergic and serotonergic neuronal subepidermal networks respectively (**Fig. S9C-D’**). To our surprise, we also detected serotonin within the germ cell cysts, (**Fig. 7A-B**) as well as genes encoding for neuromodulators and neuropeptides including *tph*, *chat*, prolactin-releasing peptide (*prrp*), luqin (*lqp*) and uncharacterized neuropeptide precursor (*np25*) (**Fig. S9A**). The widespread presence of neuromodulators within the germ cell cysts prompted us to further assess the link between ovarian neurons and germ cell cysts (**Fig. 7C**). Importantly, most of the ligand/receptor pairs used for this analysis have been previously identified and some even de-orphanized in sea stars and sea urchins (74–80). We found that neuropeptides involved in the function and regulation of the mammalian HPG axis, including gonadotropin-releasing hormone (*gnrh*), kisspeptin (*kpp*), *oxy/vp* (81) and the echinoderm muscle relaxant neuropeptide F-Salmf (76) were specifically expressed in the neuron cell type group and in both germ cell groups (**Fig. 7C**). Their receptors were predominantly expressed in the germ cell cyst clusters, suggesting crosstalk between neurons and germ cell cysts (**Fig. S7C**). Similarly, the receptors for serotonin, dopamine and GABA were found expressed within the germ cell cyst clusters (**Fig. S9B**). On the other hand, we found the ngffyamide (*ngffap*) neuropeptide to be expressed primarily in the germ and oocytes cell type groups and not in the neurons, while the latter contained transcripts for its receptor (**Fig. S7C**). Moreover, we also found neuropeptides including *prrp* and *lqp* as well as their receptors expressed by both germ cell cysts and neurons (**Fig. S7C**). Last, we report additional putative neuropeptidergic crosstalk of the ovarian neurons with the muscles and the ovarian epithelium (**Fig. S7C**).

**Figure 7.**
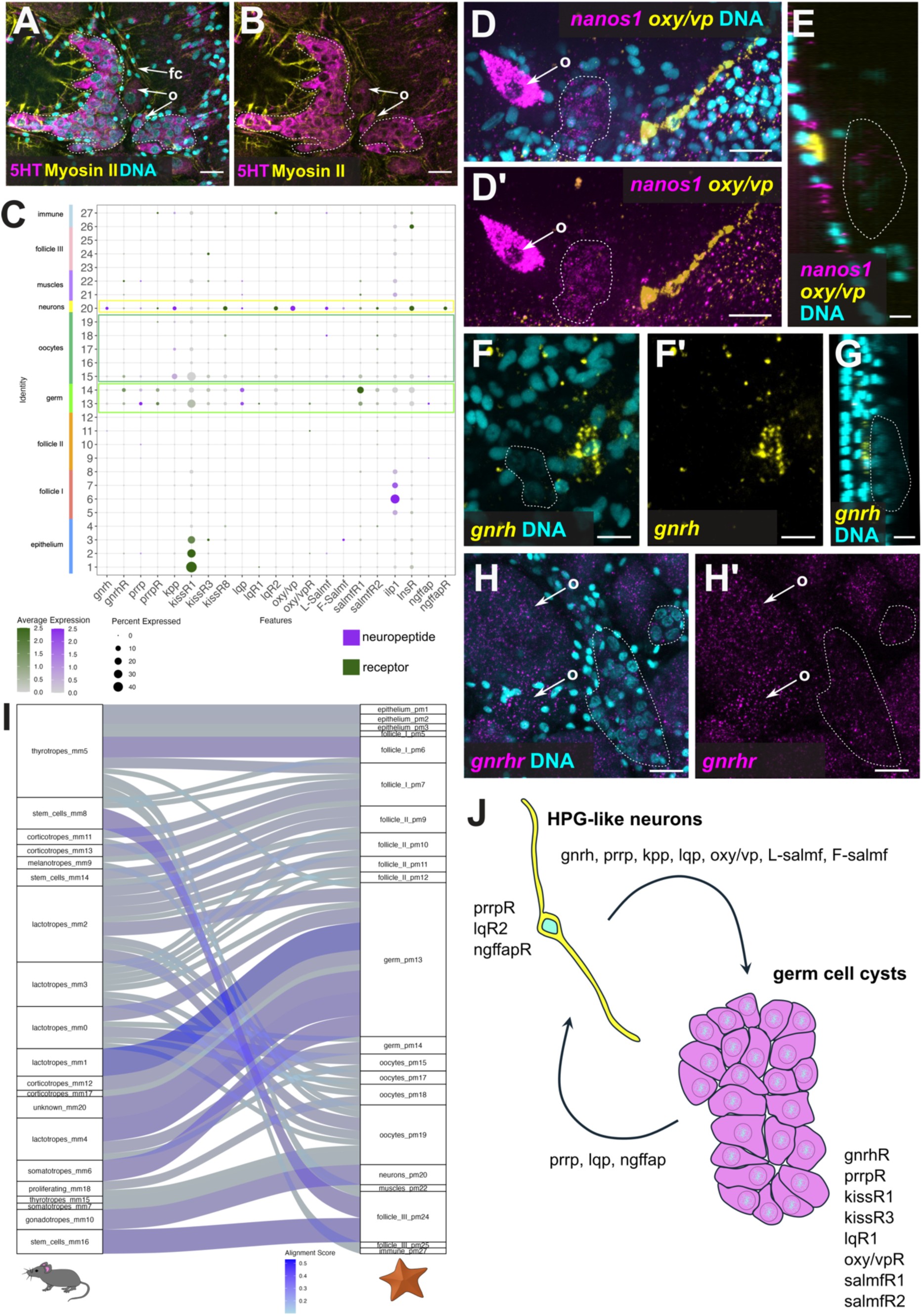
Neuropeptidergic profile of the *P. miniata* intrinsic ovarian nervous system. **(A-B)** Immunofluorescence of 5HT and Myosin II. Dotted lines outline the germ cell cysts. Scale bars 20 μm. **(C)** Dotplot showing the average expression of the reconstructed and/or deorphanized pairs of neuropeptides/receptors. **(D-D’)** HCR for *oxy/vp* and *nanos1* shows the relatively short distance between the oxy/vp positive neurons and the germ cell cysts. Scale bars in panels are 20 μm. Dotted lines outline the germ cell cysts. **(E)** Orthogonal views of the *oxy/vp* and *nanos1* HCR, shows the close proximity of the *oxy/vp* neuron in respect to the *nanos1* positive germ cell cysts. Dotted lines outline the germ cell cysts. Scale bar is 10 μm. **(F-F’)** HCR for *gnrh* in subepithelial ovarian neurons. Scale bars in panels are 20 μm. Dotted lines outline the germ cell cysts. **(G)** Orthogonal views of a *gnrh* HCR, highlights the spatial organization of the *gnrh* positive neurons in close proximity to the germ cell cysts. Scale bar is 10 μm. Dotted lines outline the germ cell cysts. **(H-H’)** HCR for *gnrhr* confirms the single cell predictions of expression in the germ lineage. Scale bars are 20 μm. Dotted lines outline the germ cell cysts. **(I)** Sankey plot depicting the alignment scores between adult sea star ovarian cell atlas with the adult mouse pituitary gland one. **(J)** Schematic representation of the components participating in the predicted crosstalk between the intrinsic ovarian neurons and the germ cell cysts. Nuclei are labelled with DAPI (cyan). fc, follicle cells; o, oocytes.

To test these predictions, we examined expression of *oxy/vp* as well as for the lowly expressed *gnrh* and *gnrhr*. Using HCR, we found that oxy/vp are expressed exclusively in rare subepidermal neurons (1-2 per ovarian lobe, **Fig. 7D-D’, S9 E-E’**). These *oxy/vp* positive neurons were in close proximity to the germ cell cysts, suggesting potential crosstalk (**Fig. 7E**). Similarly, we detected *gnrh+* cells in proximity to germ cell cysts, while *gnrhr* was expressed both in the cysts and oocytes (**Fig. 7F-H’**). Previous studies in the adult sea star *A. rubens* reported that *gnrh* is expressed predominantly within the sea star’s nervous system (oral nerve ring and radial nerve cords). Consistent with this prior work, and supporting the specificity of our HCR probes, *gnrh* and *gnrhr* were in complementary regions of the oral nerve ring and the radial nerve cord (**Fig. S9F-F’**).

Taken together, our analysis found that many known hypothalamic neuromodulators involved in the vertebrate HPG axis such as 5HT, dopamine, GABA, GNRH, KPP and OXY/VP (81) as well as the pituitary gland secreted PrRP (82) are all present in the sea star ovarian neurons. To further test if such a system is in place in sea star ovaries, we assessed the similarity between sea star cell type families with the mammalian pituitary gland using SAMap (**Fig. 7I**). Surprisingly, this analysis showed that the sea star neurons exclusively align with pituitary gland gonadotropes, which upon stimulation by hypothalamic *gnrh* produce the gonadotropins that control sex hormone production, oogenesis, and ovulation in mammals (83). Our comparison showed alignments between pituitary gland thyrotropes with the sea star follicle cells (follicle I) while the germ cell cysts aligned with the pituitary gland lactotropes (**Fig. 7I**). Altogether, our results reveal a diverse and putatively multifunctional intrinsic ovarian nervous system that may communicate with the germ cell cysts (**Fig. 7J**). Lastly, our data suggest that the sea star ovarian nervous system has a dual hypothalamic-like and gonadotropic-like state that could act as a localized HPG-like center operating in parallel or independently from the oral nerve ring and radial nerve cords to regulate sea star oogenesis, oocyte maturation and spawning behavior.

## Discussion

Sexual reproduction depends on the interactions between multiple cell types within the ovary. Ovarian structure varies across animals, reflecting differences in reproductive strategies and adaptations to different environments. For instance, many marine invertebrates follow seasonal patterns and release large quantities of gametes into the water column, whereas in mammals seasonal mating patterns result in the fertilization of only a limited number of oocytes within the oviduct. Despite these differences, in most animals studied, the ovary serves two critical roles as both an endocrine organ that produces steroid hormones, and as the niche for oocyte development. Here we took advantage of the easily accessible, optically transparent and highly fecund sea star ovary, representing an extreme case of reproductive output, to explore the identity and the evolutionary origins of the ovarian cell types in a non-chordate deuterostome. Our single cell atlas identified nine cell type groups distributed across twenty-seven transcriptionally distinct clusters which we then spatially mapped within the ovary. We show that the ovary is made of seven somatic cell groups corresponding to ciliated epithelium, neurons, muscles, immune cells and three different types of follicle cells, corroborating previous morphological studies (36). We also found cell groups corresponding to young oocytes as well as proliferating germ cells organized into cysts. Single cell trajectory analysis finds connections between the cysts and the oocytes and shows how the gene expression changes during early oogenesis. Collectively, this analysis provides a framework for understanding echinoderm oogenesis at a new depth, and a valuable resource for future investigation of reproductive biology and cell type evolution.

### Sea star follicle cells are a heterotypic group

Follicle cells that enwrap oocytes serve important roles in oogenesis across animals (84). In insects, different types of follicle cells produce yolk components or contribute to the formation of egg protective layers (85). In mammals and fish, granulosa cells are involved in the production of estrogen, nutrient transfer and regulation of oocyte maturation (1,86,87). Theca cells instead populate the outer layers of the follicle and produce steroid hormones that are taken up and converted to estrogen by granulosa cells. Similarly in echinoderms, follicle cells produce maturation-inducing molecules, such as 1-methyladenine in sea stars, upon stimulation by the gonad stimulating substance (GSS) from the nervous system that triggers the resumption of meiosis in arrested oocytes (88,89). However, the molecular signature of follicle cells has not been well defined. Here we provide evidence for three diversified sea star follicle cell groups, one that appears to be echinoderm specific (follicle II), involved primarily in vitellogenesis, and two that express genes typically found in granulosa (follicle I) and theca cells (follicle III) in mammals. Surprisingly, our results suggest possible pre-chordate evolutionary origins for theca and granulosa cells.

In support of this hypothesis, our transcriptomic and spatial analysis discovered that several genes associated with mammalian granulosa cells are also expressed by a subset of sea star follicle cells. This includes the transcription factors *foxl2* and *gata4*, and the protein hormone inhibin beta (*inhbb*) (1,44). The forkhead transcription factor *foxl2* is a crucial member of the GRN controlling the granulosa cell proliferation and differentiation in mammals (43), and in sea stars we find it expressed by a subset of follicle cells that are in direct contact with the developing oocytes. In addition, we detected primary cilia on the surface of the follicle cells enwrapping oocytes, which is a conserved feature of estrogen-producing granulosa cells of antral follicles in mice (90). Our cross-species analysis revealed that a granulosa-like cell cluster (follicle I) consistently aligned with granulosa cell clusters in humans and mice, and with follicle cells in the rest of the species analyzed. This raises the possibility that sea star granulosa cells influence follicle development through similar mechanisms as in mammals. For instance, in mammals granulosa cells secrete several Tgfβ family ligands that influence oocyte development (91). Similarly, we find that such ligands are expressed by the granulosa-like (follicle) cells. Last, our scRNA-seq data suggest that sea star granulosa-like cells produce and respond to RA signaling, which prior work has been found to regulate granulosa cell proliferation and steroidogenesis in mammals (92). Moreover, our data show that the follicle III cell group expresses star and *cyp17a1*, genes essential for steroidogenesis in mammalian theca cells. The sea star ovarian epithelium shares similarities with theca cells in zebrafish, mouse, and fetal human ovaries, with two clusters expressing *cyp17a1* and one expressing star, indicating a potential role in steroid hormone production similar to what was previously reported in humans (93). Taken together our data suggest that the sea star ovary harbors follicle cell types evolutionarily conserved to distinct follicle cell populations in other deuterostomes as well as protostomes.

### Molecular identification of putative oogonial stem cells

An evolutionarily conserved feature of oogenesis is the formation of germ cell cysts. Clusters of cytoplasmically interconnected oogonia have been found in the ovaries of a variety of animals including insects, amphibians, reptiles, birds, and mammals (35,94–99). Here we define the germline identity of sea star germ cell nests and provide evidence for intercellular bridges within them, thus suggesting that germ cell cysts are conserved in non-chordate deuterostomes. Using 3D segmentation and reconstruction, we also defined the spatial organization of oogenesis, in which germ cell cysts are localized to the periphery of the ovarian lobe along with young oocytes organized in primary follicles. Cyst expressed genes include the conserved germline markers *piwi1*, *nanos1*, *nanos3*, *boll*, *msl3*, meiotic recombination components *sycp1* and *mlh1*, and transcriptional regulators including *figla*, *atrx1* and *cxxc1* (59,100–102). We also detect cell proliferation within the cysts, supporting them as the likely source of new oogonia and oocytes in the adult ovary (103). The conserved germline RNA helicase *ddx4/vasa* is inactive in the cysts, but is expressed in early follicles, similar to what was shown in marsupial fetal germ cells (104–106). This repression may contribute to maintaining an undifferentiated state, as seen in fruit flies where *ddx4* knockout led to arrested oocyte development (107). Additionally, we hypothesize that the self-renewal-meiotic entry decision could be regulated by RA, Wnt, Tgfβ and Hh pathways, whose respective receptors and intracellular components were found expressed in the germ cell cysts. These results support an ancient role for these pathways in oogenesis, as demonstrated in other species including mice and amphibians (108–111).

### Neuronal control of oogenesis predates the establishment of chordate HPG axis

Oogenesis and ovulation are regulated by neuroendocrine signals from the nervous system in many animals. In vertebrates, this signaling occurs between the hypothalamus, the pituitary gland, and the gonad, also known as the HPG axis. HPG signaling molecules include neuropeptides *gnrh* and *oxy/vp* that are secreted by the hypothalamus and received by gonadotropin cells within the pituitary gland. This induces the secretion of the gonadotropin hormones follicle stimulating hormone (FSH) and luteinizing hormone (LH), that in turn regulate gametogenesis within the gonad through activation of the steroidogenic pathway. In mammals, the secretion of upstream hypothalamic neuropeptides is regulated by several neuromodulators including 5HT, dopamine, GABA, PrRP and KPP (112–116). How might oogenesis and ovulation be regulated in animals like sea stars that lack a central nervous system?

Foundational work in sea stars found that soluble signals from radial nerves induce oocyte maturation and spawning, indicating that oogenesis is under neural control even in invertebrates (117). Our data further argue that the sea star ovary itself contains an intrinsic neurochemical regulatory system. For instance, we found that the epithelium, germ cells, and oocytes produce previously identified neuropeptides such as *rgpp* and *trh*, both of which have been found to be produced by the radial nerve cords and regulate oogenesis in echinoderms (118,119). We also found numerous other HPG-associated signaling factors, including 5HT, dopamine, GABA, KPP, GNRH, PrRP and OXY/VP to be expressed in the intrinsic ovarian nerve plexus and their respective receptors in the germ cell cysts. Strikingly, our data suggest the presence of a novel echinoderm ovarian cell type that features a dual molecular identity resembling chordate hypothalamic neurons and pituitary gonadotropic endocrine cells. In support of this is the similarity of the sea star neuronal cluster with the one corresponding to the mouse pituitary gland gonadotropes.

Taken together our analysis provides molecular evidence towards the presence of a hypothalamic-like and gonadotropic-like intrinsic ovarian neuroendocrine signaling system in non-chordate deuterostomes. We hypothesize that the echinoderm ovarian neurons are analogous to the mammalian HPG axis elements. We speculate that during animal evolution, oogenesis was initially regulated by intrinsic neuroendocrine systems within the ovary and as organisms became more complex the regulatory role was co-opted by distinct organs such as complex interconnected nervous systems and endocrine glands.

## Acknowledgements

We thank Jamie MacKinnon, Akshay Kane, Beverly Naigles and Karen Velez, current and past members of the Swartz lab, for advice and fruitful discussions throughout the duration of this project. We thank Maria Ina Arnone (Stazione Zoologica Anton Dohrn) for helping by providing the 1e11 and Mhc antibodies, and *S. purpuratus* ovaries. We also thank Rossella Annunziata (Stazione Zoologica Anton Dohrn) for kindly providing us with *H. tubulosa* ovaries. We would also like to thank Louie Kerr, the Central Microscopy Facility at the Marine Biological Laboratory (Woods Hole, MA, USA) and Chris Bjornsson (ZEISS), for imaging assistance with confocal microscopy. We are grateful to Lisa Abbo and the Marine Resources Center staff for providing excellent animal care and husbandry of adult *P. miniata* used in this work.

## Author contributions

Conceptualization: P.P., S.Z.S.; Methodology: P.P., S.Z.S.; Formal analysis: P.P., C.W., D.V.; Investigation: P.P., S.Z.S.; Writing - original draft: P.P.; Writing - review & editing: P.P., C.W., D.V., S.Z.S.; Visualization: P.P., C.W.; Supervision: S.Z.S.; Project administration: S.Z.S.; Funding acquisition: S.Z.S.

## Supplementary Figures

**Figure S1.**
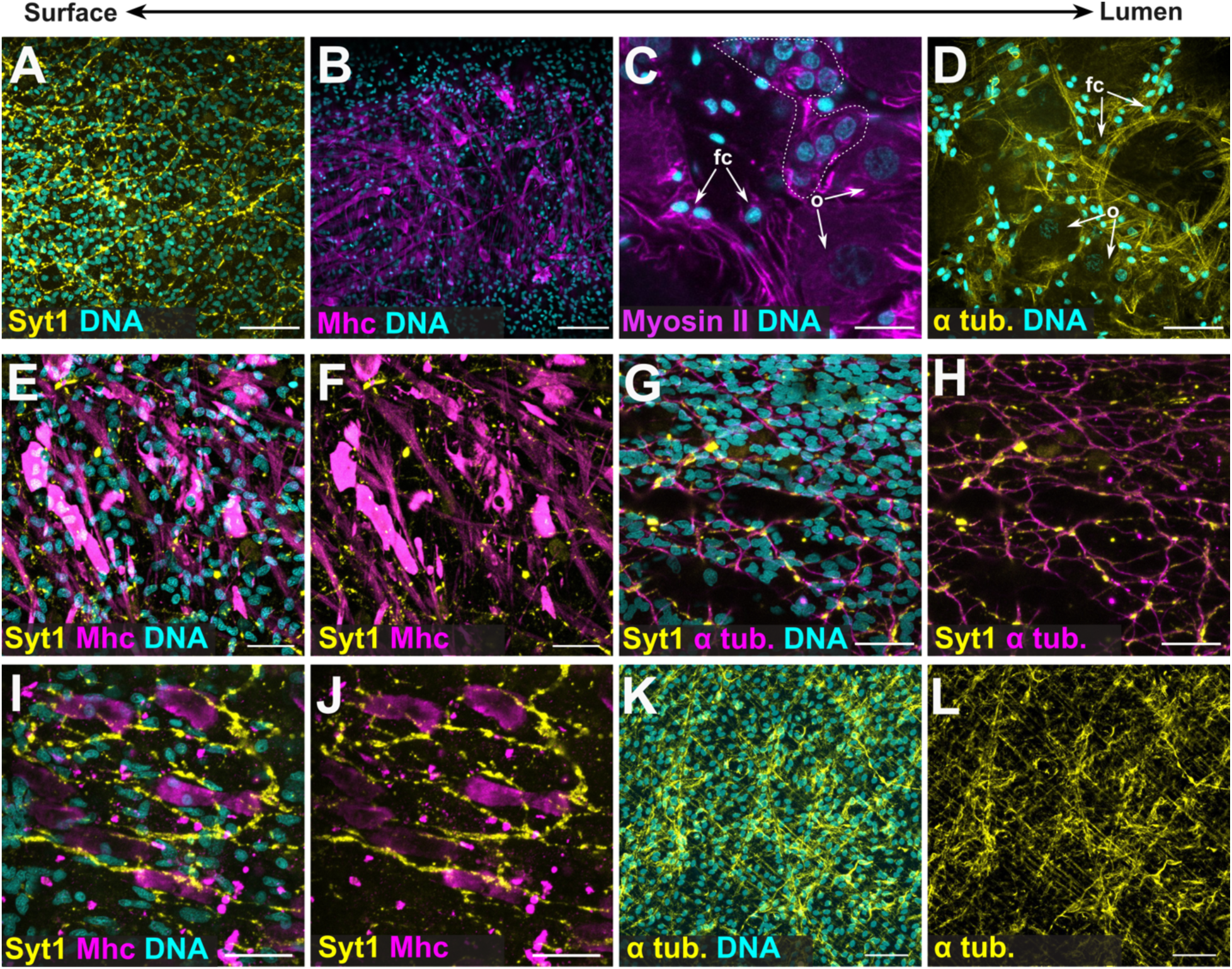
Immunohistochemical identification of echinoderm ovarian cell types. **(A-D)** IHC for Syt1 labeling neurons (A), for Mhc labeling muscles (B) and for myosin (C) and α-tubulin (D) labeling follicle cells. Scale bars for panels A and D are 40 μm; for panel B 60 μm; for C 20 μm. **(E-F)** IHC for Syt1 and Mhc to visualize the interconnection of ovarian neurons and muscles. Scale bars for panels are 20 μm. **(G-H)** IHC for Syt1 and α-tubulin co-labels the ovarian neurons. Scale bars for panels are 20 μm. **(I-J)** IHC for Syt1 and Mhc confirms the presence of the neurons and muscles respectively in the sea urchin *S. purpuratus* ovary. Scale bars are 25 μm. **(K-L)** IHC for α tubulin labels the ovarian neurons in the sea cucumber *H. tubulosa* ovary. Scale bars are 50 μm. Nuclei were labelled with DAPI (cyan). fc, follicle cells; o, oocytes.

**Figure S2.**
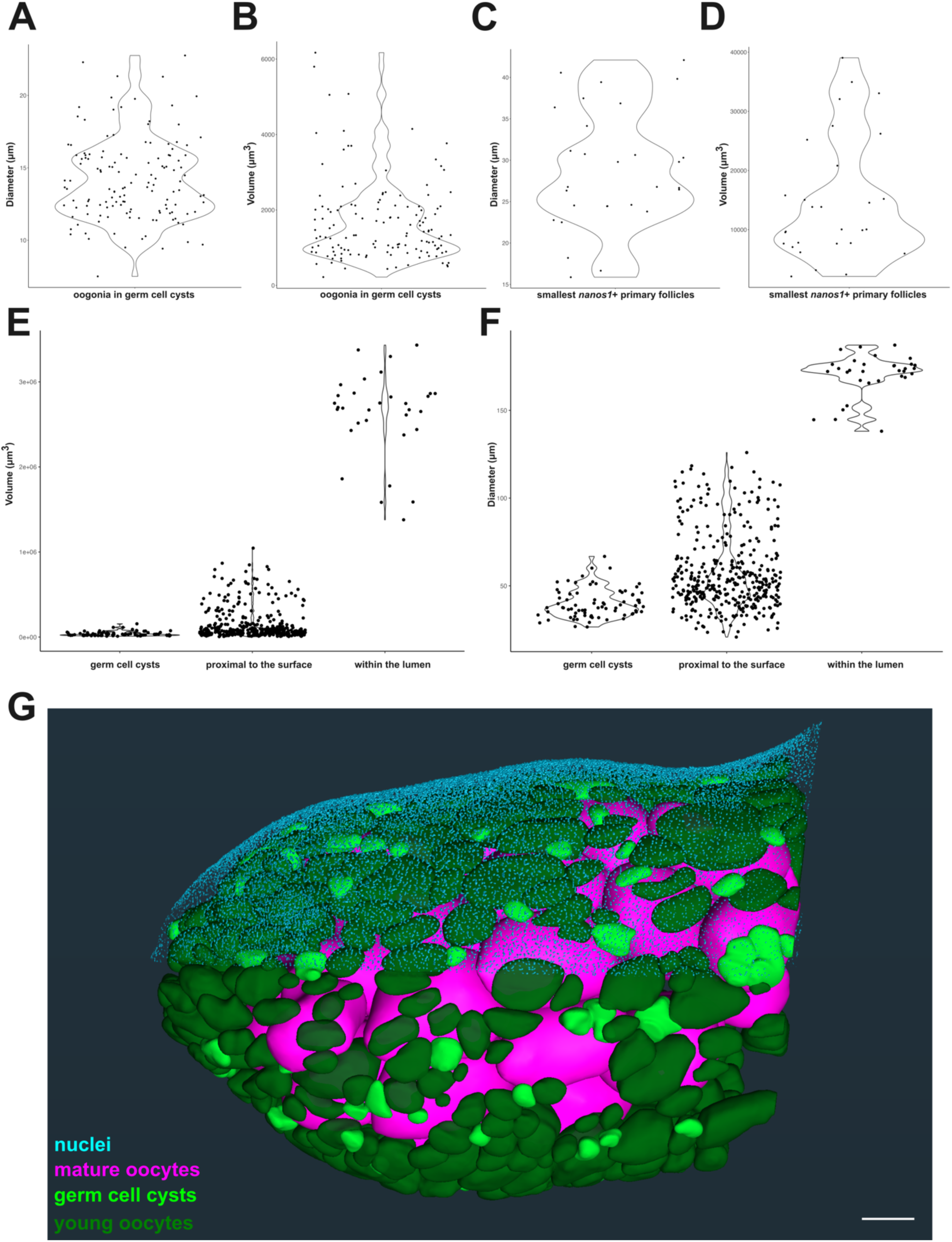
Size and spatial distributions of oogonia and oocytes in the sea star ovary. **(A-B)** Violin plot showing the experimentally measured diameter (A) and estimated volume (B) of each oogonium within germ cell cysts. **(C-D)** Violin plot showing the measured diameter (C) and estimated volume (D) of the smallest *nanos1* positive oocyte in close proximity to the germ cell cysts. **(E-F)** Violin plot of the segmentation-based calculated volumes (E) and estimated diameter (F) of germ cell cysts, oocytes close to the ovarian surface, and those within the lumen. **(G)** 3D rendering of segmented germ cell cysts and oocytes. Scale bar is 100 μm. Ovary seen from a side view. Estimated volumes or diameters were calculated based on experimentally measured diameters or volumes respectively, assuming the shape of the oogonia and oocytes is spherical.

**Figure S3.**
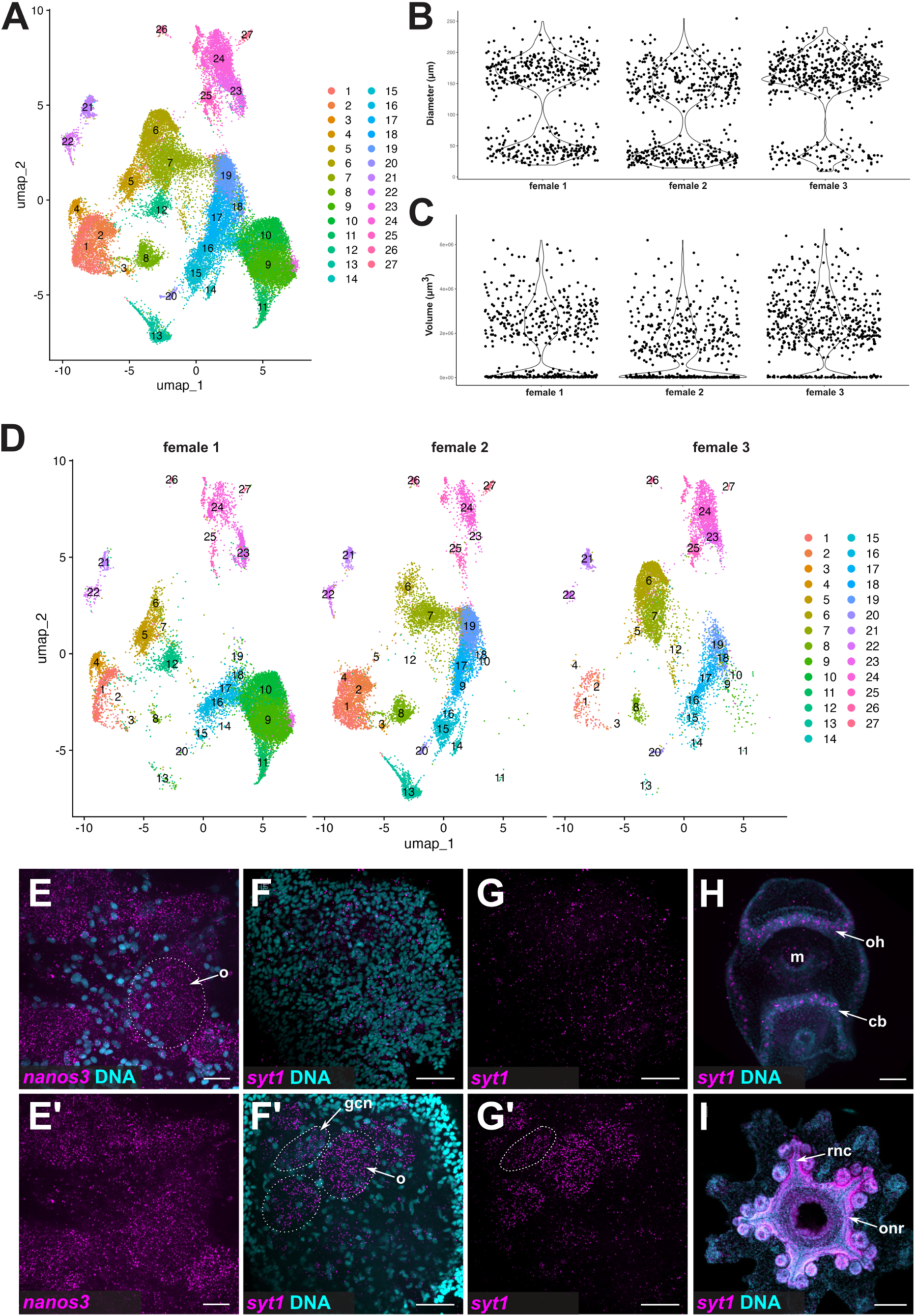
*P. miniata* ovary cell atlas and cell contribution of the different libraries. **(A)** Integrated UMAP of the three different scRNA-seq libraries, in which color code is indicative of different clusters **(B-C)** Violin plot showing the experimentally measured diameter (B) and estimated volume (C) of the oogonia and oocytes within the ovaries used for the reconstruction of the single cell atlas. Due to the capturing system limitations only the oocytes with a size ≤ 40μm in diameter are included in the single cell atlas. **(D)** Integrated UMAP of the three different scRNA-seq libraries, split by the cell contribution of each library. The color code is indicative of different clusters. **(E-E’)** HCR for the *nanos3* gene with (E) and without (E’) the nuclei channel (DAPI) confirmed its localization in the germ lineage. Dotted circles in panel E are indicative of oocytes. Scale bars are 20 μm. **(F-G’)** HCR for the *syt1* gene with (F-F’) and without (G-G’) the nuclei channel (DAPI) shows expression in the neurons (F-G) as well as the germ cell cysts and oocytes (F’-G’). Dotted circles in F’ are indicative of oocytes. Germ cell cysts in panels F’ and G’ are outlined with dotted lines. Scale bars are 20 μm. **(H-I) HCR** for the *syt1* gene with (G) and without (G’) the nuclei channel (DAPI) shows expression of the gene only in the nervous system of the *P. miniata* larvae (H) and post-metamorphic juveniles (I), confirming the specificity of the probe. Scale bars for panels H and I are 50 and 100 μm respectively. cb, ciliary band; m, mouth, oocytes; oh, oral hood; onr, oral nerve ring; rnc, radial nerve cord.

**Figure S4.**
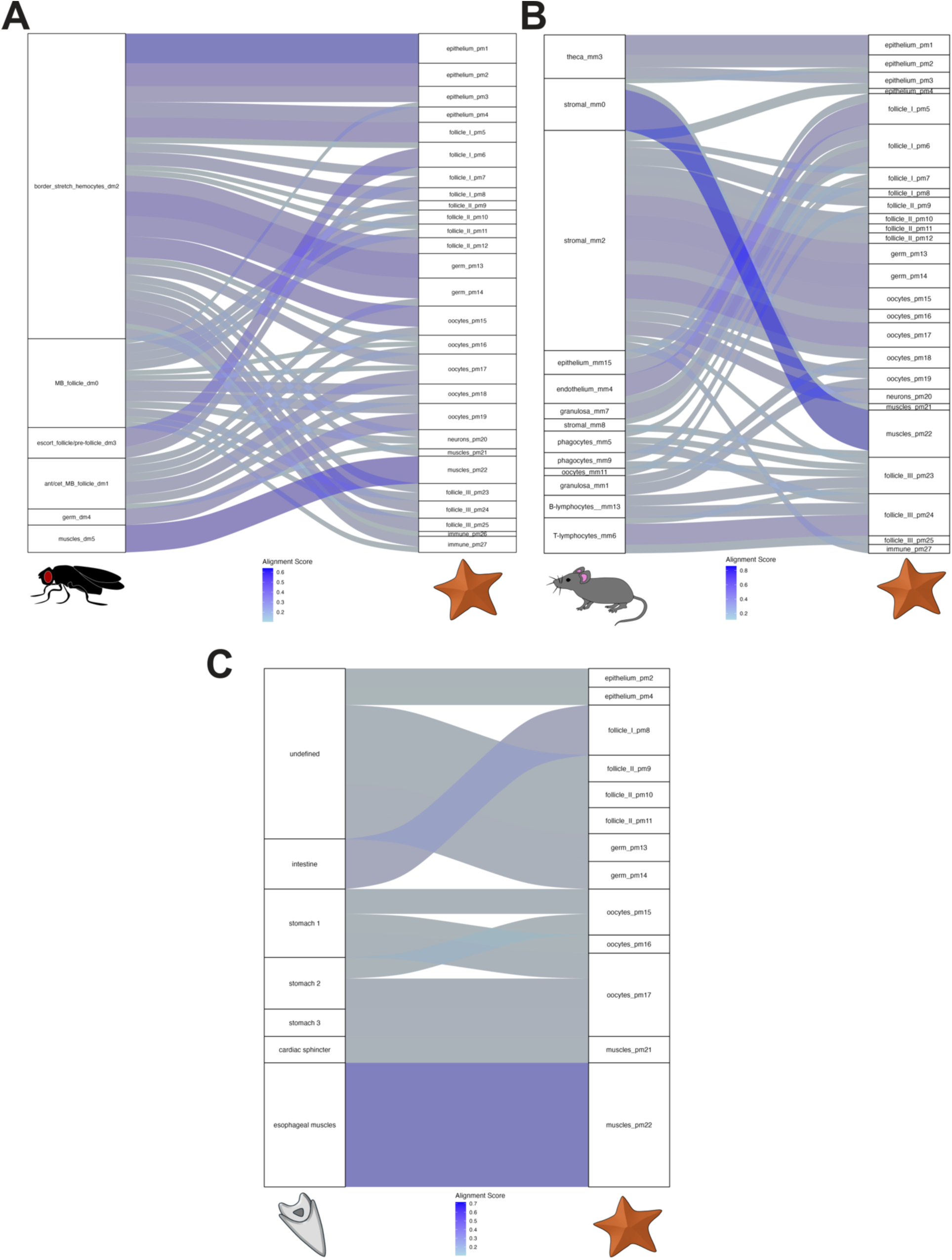
Cross-species comparison of sea star ovarian cell types with protostome and deuterostome representatives with SAMap. Sankey plot depicting the alignment scores between adult sea star ovarian cell atlas with the adult fruit fly ovary **(A)**, adult mouse ovary **(B)** and with sea urchin larva **(C)**. Alignment scores equal or greater to 0.1 were considered significant.

**Figure S5.**
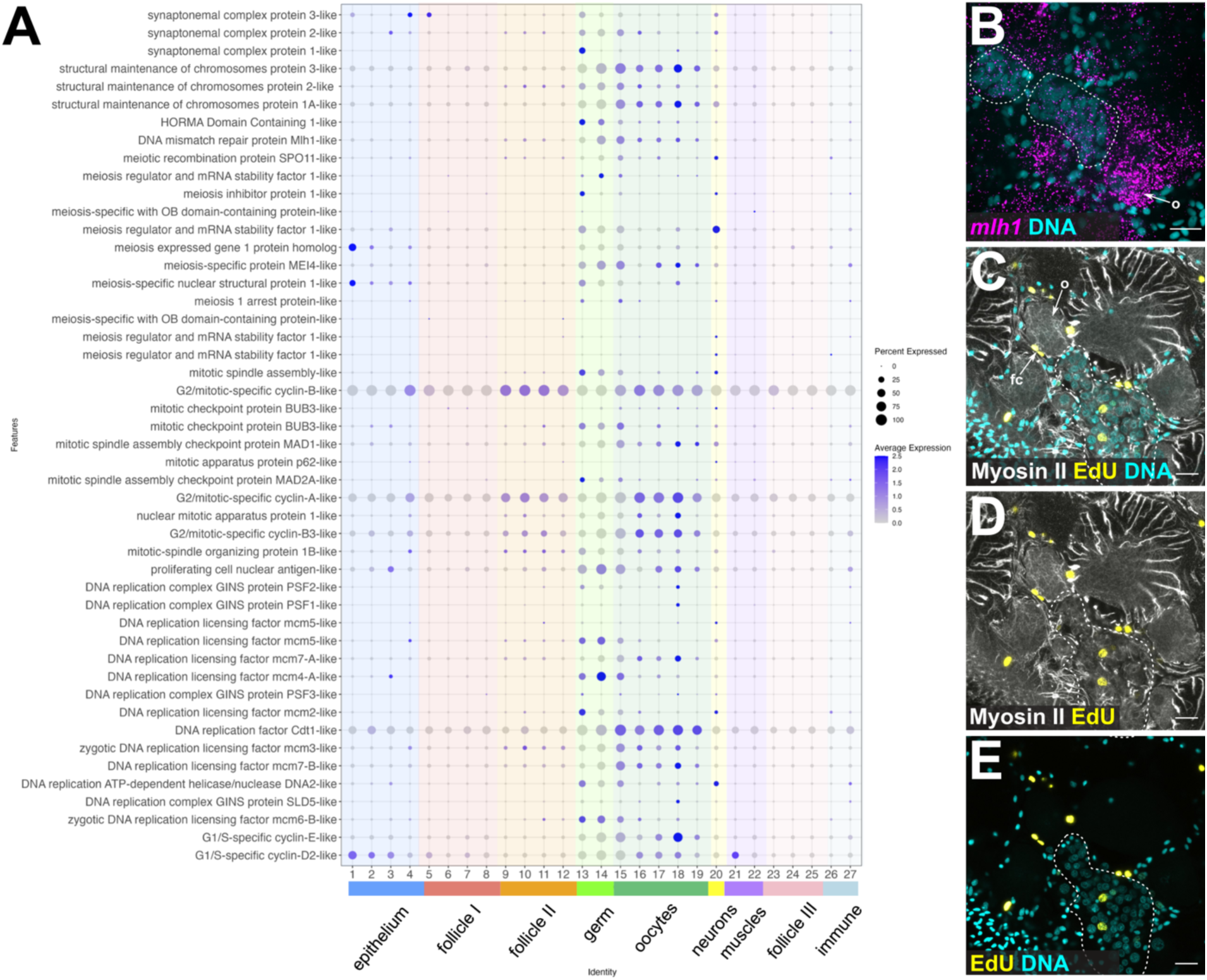
Identification of mitotic and meiotic molecular signatures in the sea star ovary. **(A)** Dotplot showing the average expression of the genes used for the scoring analysis shown in Fig.4 A. **(B)** HCR for the meiotic marker *mlh1* labelling the germ cell cysts and oocytes of various sizes. **(C-E)** IHC for Myosin II labeling cell peripheries with the EdU proliferation assay, confirms that cells from both the somatic and germ lineages are actively proliferating in the adult sea star ovary. Nuclei are labelled with DAPI (cyan). Dotted lines outline the germ cell cyst in panels C-E. Scale bars are 20 μm. fc, follicle cells; o, oocytes.

**Figure S6.**
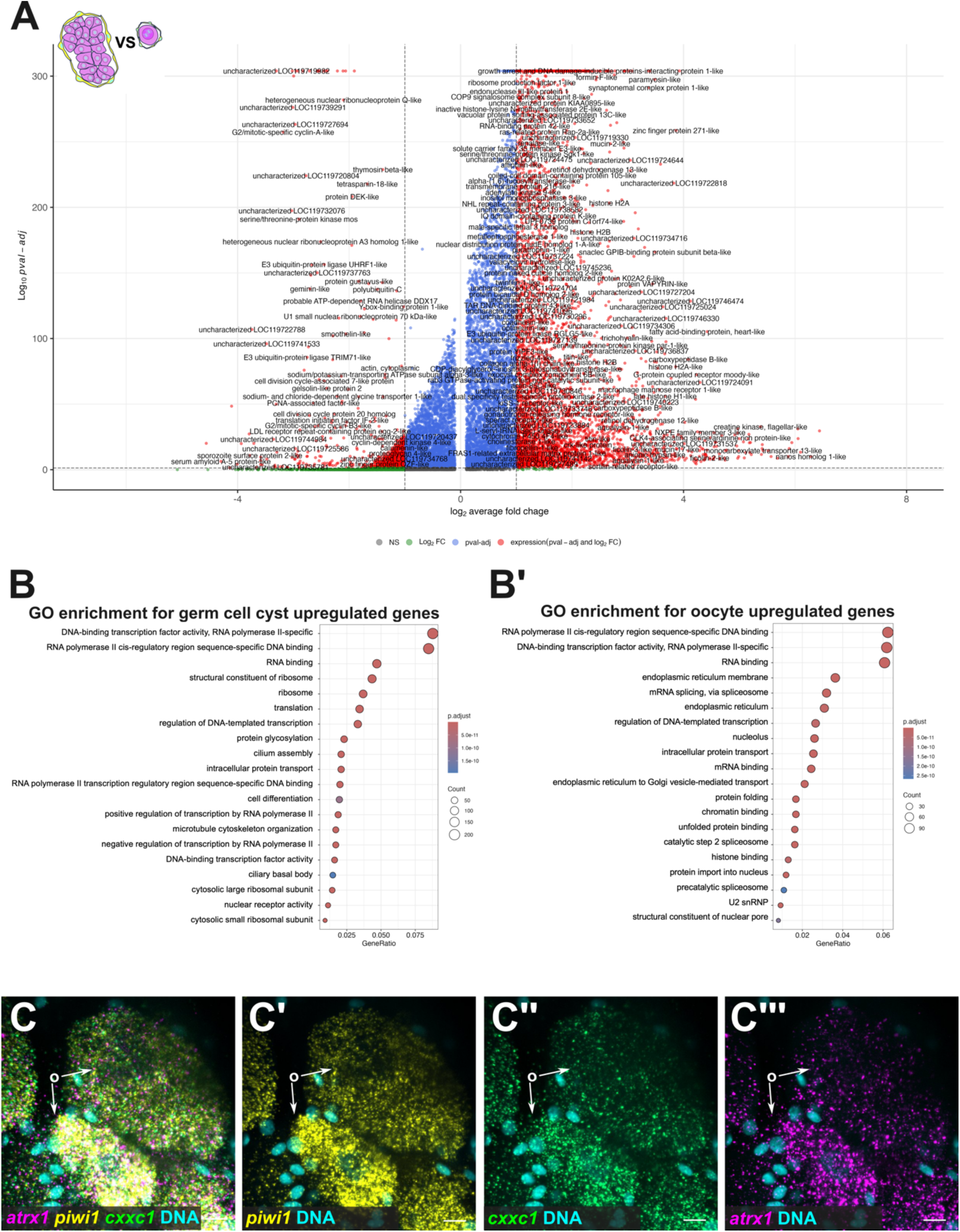
Differential expression patterns along the sea star oogenesis developmental trajectory. **(A)** Volcano plot showing the differential expression of the germ cell cyst (clusters 13 and 14) genes versus the oocyte reference group (clusters 15-19). **(B-B’)** Dotplot showing the top 20 enriched gene ontology terms associated with the germ cell cyst (B) and the oocyte (B’) upregulated genes. **(C-C’’’)** HCR for the transcriptional regulators *atrx1* and *cxxc1* paired with *piwi1* shows expression in oocytes. Scale bars are 10 μm.

**Figure S7.**
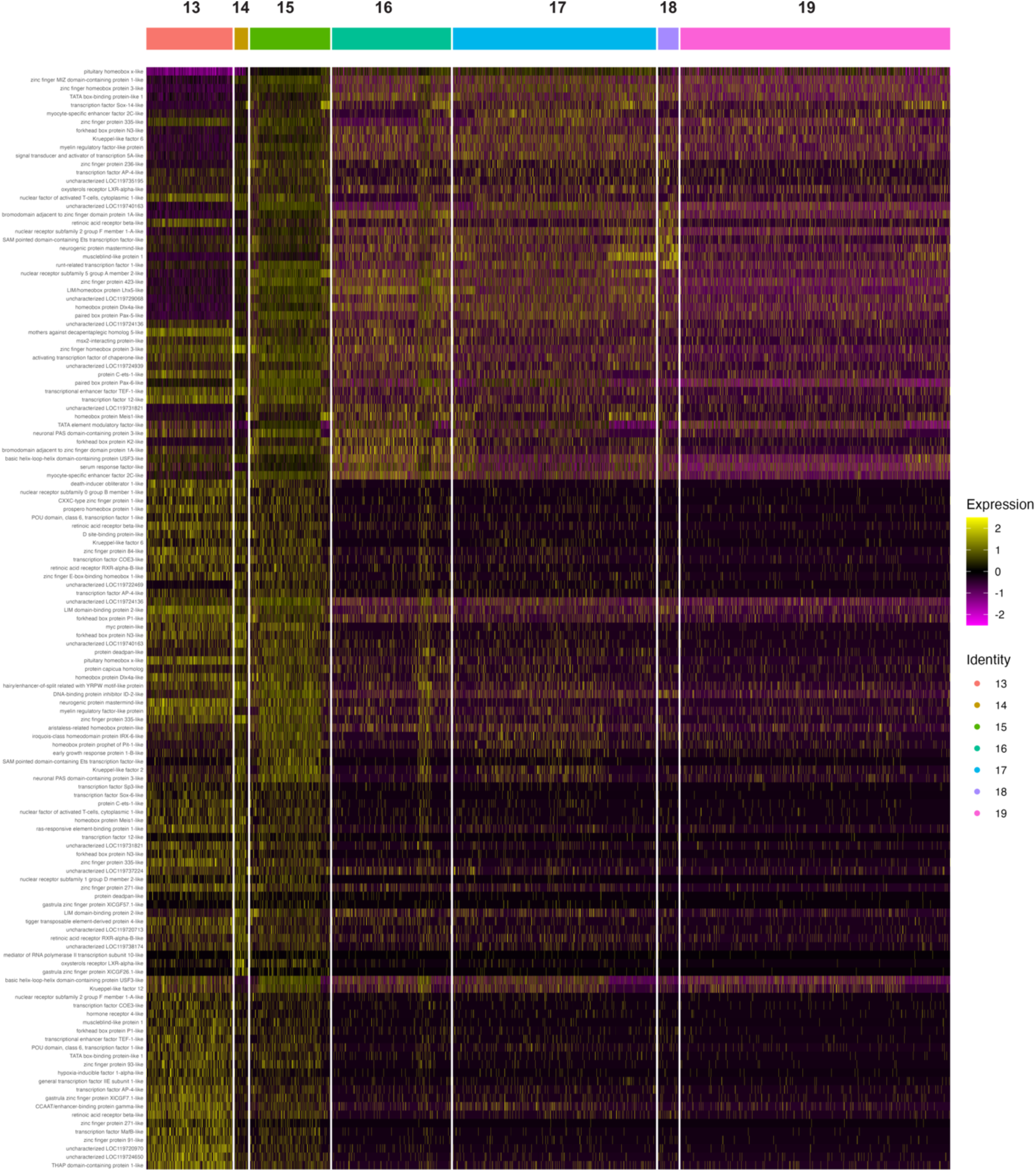
Distinct transcription factor profiles during early oogenesis. Heatmap of the top 50 differentially expressed transcription factors across the germline clusters.

**Figure S8.**
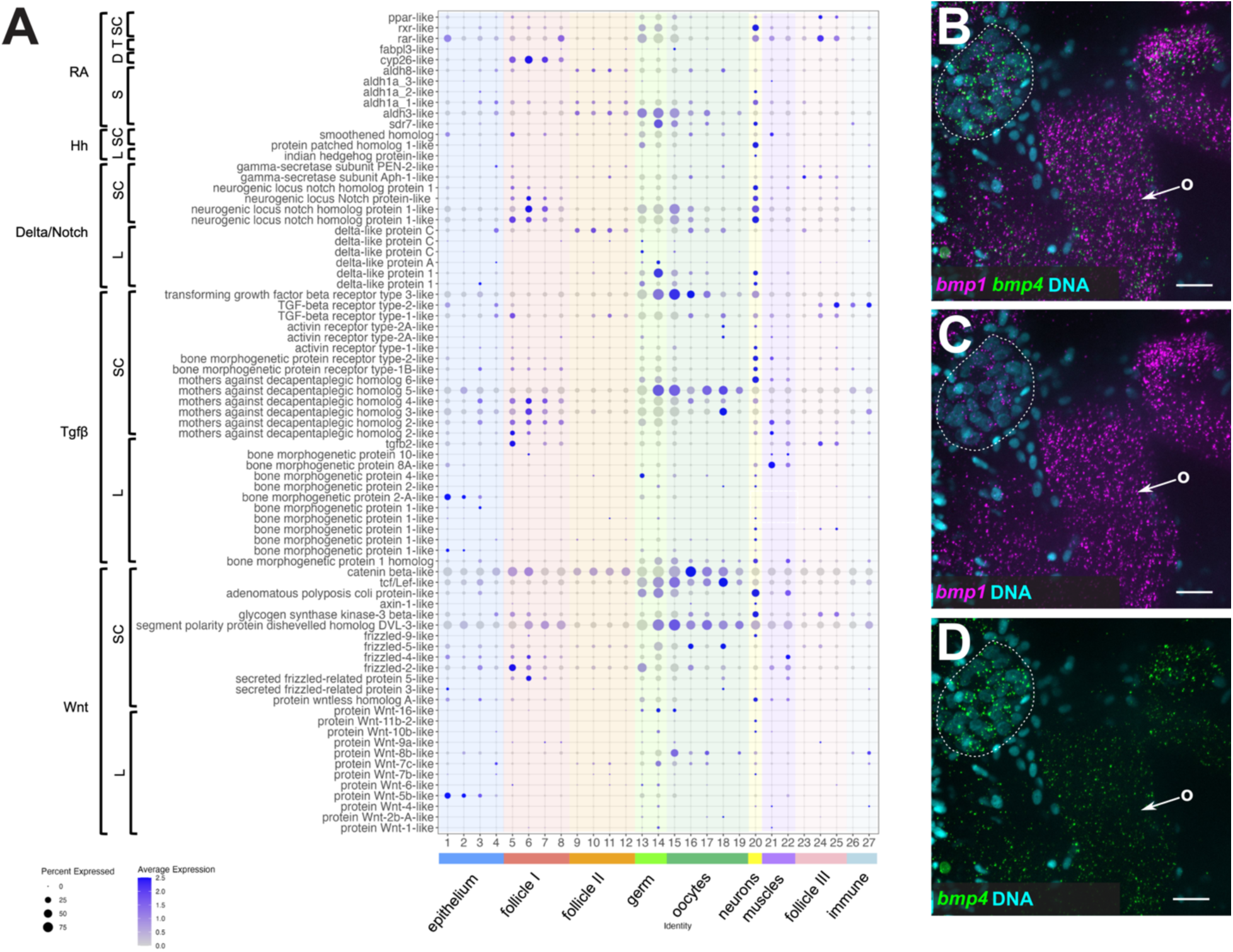
Expression patterns of evolutionarily conserved ovarian signaling pathways. **(A)** Dotplot showing the average expression of the genes used for the scoring analysis shown in Fig. 6A. D, degradation; L, ligand; S, synthesis; SC, signaling cascade; T, transport. **(B-D)** HCR for the *bmp1* and *bmp4* ligands confirms their spatial expression in the germ cell cysts and oocytes. Nuclei are labelled with DAPI (cyan). Dotted lines outline the germ cell cyst. Scale bars are 20 μm. o, oocytes.

**Figure S9.**
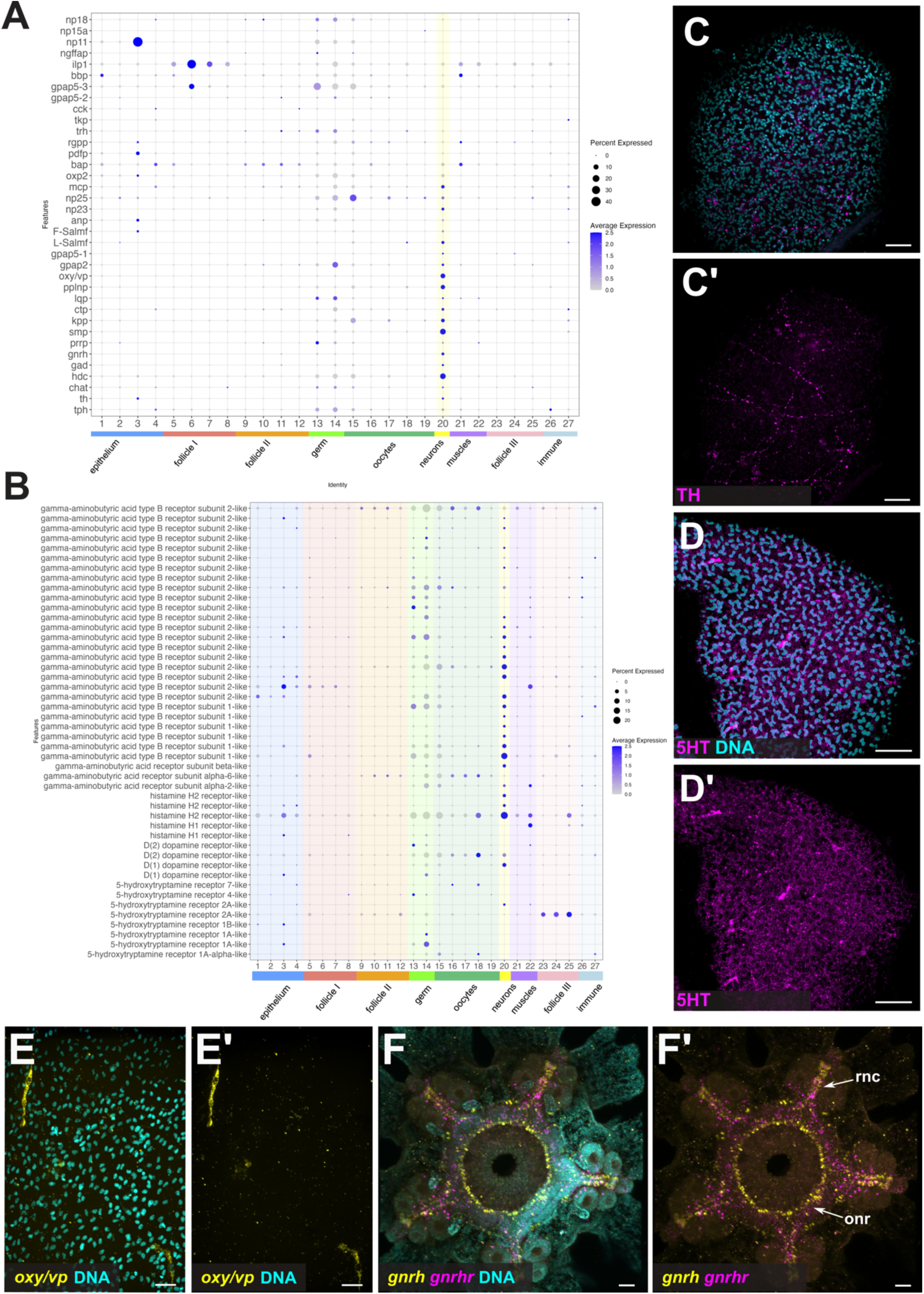
Neuropeptides and neurotransmitters expressed in the intrinsic ovarian nervous system. **(A)** Dotplot showing the average expression of known echinoderm neurotransmitter synthesizing enzymes and neuropeptides. **(B)** Dotplot showing the average expression of sea star neurotransmitter receptors. **(C-C’)** Immunofluorescence of TH shows the presence of subepidermal ovarian dopaminergic neurons. Scale bars are 40 μm. **(D-D’)** Immunofluorescence of 5HT confirms the presence of subepidermal ovarian serotonergic neurons. Scale bars are 40 μm. **(E-E’)** HCR for *oxy/vp* neuropeptide marks two *oxy/vp* positive neurons per ovarian lobe. Scale bars are 20 μm. **(F-F’)** HCR for *gnrh* and *gnrhr* shows expression in the post-metamorphic *P. miniata* nervous system confirming the specificity of the probes. Scale bars are 50 μm. onr, oral nerve ring; rnc, radial nerve cord.

## References

1. Prasasya RD, Mayo KE. Regulation of follicle formation and development by ovarian signaling pathways. In: The Ovary. Elsevier; 2019. p. 23–49.

2. Orisaka M, Tajima K, Tsang BK, Kotsuji F. Oocyte-granulosa-theca cell interactions during preantral follicular development. J Ovarian Res. 2009 Jul 9;2(1):9.

3. Tsukamura H. Kobayashi Award 2019: The neuroendocrine regulation of the mammalian reproduction. Gen Comp Endocrinol. 2022 Jan 1;315(113755):113755.

4. Grive KJ, Freiman RN. The developmental origins of the mammalian ovarian reserve. Development. 2015 Aug 1;142(15):2554–63.

5. Packer C, Tatar M, Collins A. Reproductive cessation in female mammals. Nature. 1998 Apr 23;392(6678):807–11.

6. Armstrong AR, Drummond-Barbosa D. Insulin signaling acts in adult adipocytes via GSK-3β and independently of FOXO to control Drosophila female germline stem cell numbers. Dev Biol. 2018 Aug 1;440(1):31–9.

7. Juliano CE, Swartz SZ, Wessel GM. A conserved germline multipotency program. Development. 2010 Dec;137(24):4113–26.

8. Arnone MI, Byrne M, Martinez P. Echinodermata. In: Evolutionary Developmental Biology of Invertebrates 6. Vienna: Springer Vienna; 2015. p. 1–58.

9. Bodnar AG. Marine invertebrates as models for aging research. Exp Gerontol. 2009 Aug;44(8):477–84.

10. Ernst SG. Offerings from an urchin. Dev Biol. 2011 Oct 15;358(2):285–94.

11. Arnone MI, Andrikou C, Annunziata R. Echinoderm systems for gene regulatory studies in evolution and development. Curr Opin Genet Dev. 2016 Aug;39:129–37.

12. Swartz SZ, McKay LS, Su KC, Bury L, Padeganeh A, Maddox PS, et al. Quiescent cells actively replenish CENP-A nucleosomes to maintain centromere identity and proliferative potential. Dev Cell. 2019 Oct 7;51(1):35–48.e7.

13. Paganos P, Voronov D, Musser JM, Arendt D, Arnone MI. Single-cell RNA sequencing of the Strongylocentrotus purpuratus larva reveals the blueprint of major cell types and nervous system of a non-chordate deuterostome. Elife. 2021 Nov 25;10.

14. Magri MS, Voronov D, Foley S, Martínez-García PM, Franke M, Cary GA, et al. Deep conservation of cis-regulatory elements and chromatin organization in echinoderms uncover ancestral regulatory features of animal genomes. bioRxiv. 2024.

15. Dainat J, Hereñú D, Murray KD, Davis E, Ugrin I, Crouch K, et al. NBISweden/AGAT: AGAT-v1.4.1. Zenodo. 2024.

16. Zheng GXY, Terry JM, Belgrader P, Ryvkin P, Bent ZW, Wilson R, et al. Massively parallel digital transcriptional profiling of single cells. Nat Commun. 2017 Jan 16;8(1):14049.

17. Hao Y, Stuart T, Kowalski MH, Choudhary S, Hoffman P, Hartman A, et al. Dictionary learning for integrative, multimodal and scalable single-cell analysis. Nat Biotechnol. 2024 Feb;42(2):293–304.

18. Chitiashvili T, Dror I, Kim R, Hsu FM, Chaudhari R, Pandolfi E, et al. Female human primordial germ cells display X-chromosome dosage compensation despite the absence of X-inactivation. Nat Cell Biol. 2020 Dec;22(12):1436–46.

19. Isola JVV, Ocañas SR, Hubbart CR, Ko S, Mondal SA, Hense JD, et al. A single-cell atlas of the aging mouse ovary. Nat Aging. 2024 Jan;4(1):145–62.

20. Liu Y, Kossack ME, McFaul ME, Christensen LN, Siebert S, Wyatt SR, et al. Single-cell transcriptome reveals insights into the development and function of the zebrafish ovary. Elife. 2022 May 19;11:e76014.

21. Oulhen N, Morita S, Pieplow C, Onorato TM, Foster S, Wessel G. Conservation and contrast in cell states of echinoderm ovaries. Mol Reprod Dev. 2024 Aug;91(8):e23721.

22. Rust K, Byrnes LE, Yu KS, Park JS, Sneddon JB, Tward AD, et al. A single-cell atlas and lineage analysis of the adult Drosophila ovary. Nat Commun. 2020 Nov 6;11(1):5628.

23. Allensworth-James M, Banik J, Odle A, Hardy L, Lagasse A, Moreira ARS, et al. Control of the anterior pituitary cell lineage regulator POU1F1 by the stem cell determinant Musashi. Endocrinology. 2021 Mar 1;162(3):bqaa245.

24. Tarashansky AJ, Musser JM, Khariton M, Li P, Arendt D, Quake SR, et al. Mapping single-cell atlases throughout Metazoa unravels cell type evolution. Elife. 2021 May 4;10:e66747.

25. Piovani L, Leite DJ, Yañez Guerra LA, Simpson F, Musser JM, Salvador-Martínez I, et al. Single-cell atlases of two lophotrochozoan larvae highlight their complex evolutionary histories. Sci Adv. 2023 Aug 2;9(31):eadg6034.

26. Cao J, Spielmann M, Qiu X, Huang X, Ibrahim DM, Hill AJ, et al. The single-cell transcriptional landscape of mammalian organogenesis. Nature. 2019 Feb;566(7745):496–502.

27. Hörl D, Rojas Rusak F, Preusser F, Tillberg P, Randel N, Chhetri RK, et al. BigStitcher: reconstructing high-resolution image datasets of cleared and expanded samples. Nat Methods. 2019 Sep;16(9):870–4.

28. Formery L, Peluso P, Kohnle I, Malnick J, Thompson JR, Pitel M, et al. Molecular evidence of anteroposterior patterning in adult echinoderms. Nature. 2023 Nov;623(7987):555–61.

29. Kuehn E, Clausen DS, Null RW, Metzger BM, Willis AD, Özpolat BD. Segment number threshold determines juvenile onset of germline cluster expansion in Platynereis dumerilii. J Exp Zool B Mol Dev Evol. 2022 Jun;338(4):225–40.

30. Paganos P, Caccavale F, La Vecchia C, D’Aniello E, D’Aniello S, Arnone MI. FISH for all: A fast and efficient fluorescent in situ hybridization (FISH) protocol for marine embryos and larvae. Front Physiol. 2022 Apr 19;13:878062.

31. Perillo M, Paganos P, Spurrell M, Arnone MI, Wessel GM. Methodology for whole mount and fluorescent RNA in situ hybridization in echinoderms: Single, double, and beyond. In: Methods in Molecular Biology. New York, NY: Springer US; 2021. p. 195–216. (Methods in molecular biology (Clifton, N.J.)).

32. Ćorić A, Stockinger AW, Schaffer P, Rokvić D, Tessmar-Raible K, Raible F. A fast and versatile method for simultaneous HCR, immunohistochemistry and Edu labeling (SHInE). Integr Comp Biol. 2023 Aug 23;63(2):372–81.

33. Mita M, Osugi T, Takahashi T, Watanabe T, Satake H. Mechanism of gamete shedding in starfish: Involvement of acetylcholine in extracellular Ca2+-dependent contraction of gonadal walls. Gen Comp Endocrinol. 2020 May 1;290(113401):113401.

34. Zickler D, Kleckner N. The leptotene-zygotene transition of meiosis. Annu Rev Genet. 1998;32:619–97.

35. Pepling ME. From primordial germ cell to primordial follicle: mammalian female germ cell development. Genesis. 2006 Dec;44(12):622–32.

36. Schoenmakers HJ, Colenbrander PH, Peute J, van Oordt PG. Anatomy of the ovaries of the starfish Asterias rubens (Echinodermata). A histological and ultrastructural study. Cell Tissue Res. 1981;217(3):577–97.

37. Fresques T, Swartz SZ, Juliano C, Morino Y, Kikuchi M, Akasaka K, et al. The diversity of nanos expression in echinoderm embryos supports different mechanisms in germ cell specification. Evol Dev. 2016 Jul;18(4):267–78.

38. Reich A, Dunn C, Akasaka K, Wessel G. Phylogenomic analyses of Echinodermata support the sister groups of Asterozoa and Echinozoa. PLoS One. 2015 Mar 20;10(3):e0119627.

39. Zazueta-Novoa V, Onorato TM, Reyes G, Oulhen N, Wessel GM. Complexity of yolk proteins and their dynamics in the sea star Patiria miniata. Biol Bull. 2016 Jun;230(3):209–19.

40. Xia L, Yin GL, Long Y, Sun F, Wang BB, Zhu Y, et al. Essential role of CFAP77-CCDC105-TEX43 subcomplex in connecting axonemal A and B tubules for mammalian sperm motility. Cell Biology. bioRxiv. 2025.

41. Fan X, Bialecka M, Moustakas I, Lam E, Torrens-Juaneda V, Borggreven NV, et al. Single-cell reconstruction of follicular remodeling in the human adult ovary. Nat Commun. 2019 Jul 18;10(1):3164.

42. Juengel JL, Bibby AH, Reader KL, Lun S, Quirke LD, Haydon LJ, et al. The role of transforming growth factor-beta (TGF-beta) during ovarian follicular development in sheep. Reprod Biol Endocrinol. 2004 Nov 25;2(1):78.

43. Pisarska MD, Barlow G, Kuo FT. Minireview: roles of the forkhead transcription factor FOXL2 in granulosa cell biology and pathology. Endocrinology. 2011 Apr;152(4):1199–208.

44. Bennett J, Baumgarten SC, Stocco C. GATA4 and GATA6 silencing in ovarian granulosa cells affects levels of mRNAs involved in steroidogenesis, extracellular structure organization, IGF-I activity, and apoptosis. Endocrinology. 2013 Dec;154(12):4845–58.

45. Liu Z, Castrillon DH, Zhou W, Richards JS. FOXO1/3 depletion in granulosa cells alters follicle growth, death and regulation of pituitary FSH. Mol Endocrinol. 2013 Feb;27(2):238–52.

46. Vos MC, van der Wurff AA, Last JTJ, de Boed EAM, Smeenk JMJ, van Kuppevelt TH, et al. Immunohistochemical expression of MMP-14 and MMP-2, and MMP-2 activity during human ovarian follicular development. Reprod Biol Endocrinol. 2014 Jan 31;12(1):12.

47. Abramovich D, Rodriguez Celin A, Hernandez F, Tesone M, Parborell F. Spatiotemporal analysis of the protein expression of angiogenic factors and their related receptors during folliculogenesis in rats with and without hormonal treatment. J Reprod Fertil. 2009 Feb;137(2):309–20.

48. Kiriakidou M, McAllister JM, Sugawara T, Strauss JF 3rd. Expression of steroidogenic acute regulatory protein (StAR) in the human ovary. J Clin Endocrinol Metab. 1996 Nov;81(11):4122–8.

49. Yamamoto S, Konishi I, Tsuruta Y, Nanbu K, Mandai M, Kuroda H, et al. Expression of vascular endothelial growth factor (VEGF) during folliculogenesis and corpus luteum formation in the human ovary. Gynecol Endocrinol. 1997 Dec;11(6):371–81.

50. Estienne A, Relav L, Benkoura M, Monniaux D, Morin F, Fabre S, et al. Endothelial cell-derived fibroblast growth factor-18 regulates ovarian function in sheep. J Cell Physiol. 2022 May;237(5):2528–38.

51. Chai C, Liang L, Mikkelsen NS, Wang W, Zhao W, Sun C, et al. Single-cell transcriptome analysis of epithelial, immune, and stromal signatures and interactions in human ovarian cancer. Commun Biol. 2024 Jan 26;7(1):131.

52. Bayly-Jones C, Pang SS, Spicer BA, Whisstock JC, Dunstone MA. Ancient but not forgotten: New insights into MPEG1, a macrophage perforin-like immune effector. Front Immunol. 2020 Oct 15;11:581906.

53. Bülow S, Zeller L, Werner M, Toelge M, Holzinger J, Entzian C, et al. Bactericidal/permeability-increasing protein is an enhancer of bacterial lipoprotein recognition. Front Immunol. 2018 Dec 5;9:2768.

54. Baldwin LA, Hoff JT, Lefringhouse J, Zhang M, Jia C, Liu Z, et al. CD151-α3β1 integrin complexes suppress ovarian tumor growth by repressing slug-mediated EMT and canonical Wnt signaling. Oncotarget. 2014 Dec 15;5(23):12203–17.

55. Wood NJ, Mattiello T, Rowe ML, Ward L, Perillo M, Arnone MI, et al. Neuropeptidergic systems in Pluteus larvae of the sea urchin Strongylocentrotus purpuratus: Neurochemical complexity in a “simple” nervous system. Front Endocrinol (Lausanne). 2018 Oct 25;9:628.

56. Pagowski V. A description of the bat star nervous system throughout larval ontogeny. Evol Dev. 2024 Jan;26(1):e12468.

57. Kee K, Angeles VT, Flores M, Nguyen HN, Reijo Pera RA. Human DAZL, DAZ and BOULE genes modulate primordial germ-cell and haploid gamete formation. Nature. 2009 Nov 12;462(7270):222–5.

58. Gonzalez LE, Tang X, Lin H. Maternal Piwi regulates primordial germ cell development to ensure the fertility of female progeny in Drosophila. Genetics. 2021 Aug 26;219(1).

59. McCarthy A, Sarkar K, Martin ET, Upadhyay M, Jang S, Williams ND, et al. Msl3 promotes germline stem cell differentiation in female Drosophila. Development. 2022 Jan 1;149(1).

60. Sage J, Martin L, Meuwissen R, Heyting C, Cuzin F, Rassoulzadegan M. Temporal and spatial control of the Sycp1 gene transcription in the mouse meiosis: regulatory elements active in the male are not sufficient for expression in the female gonad. Mech Dev. 1999 Jan;80(1):29–39.

61. Hu W, Gauthier L, Baibakov B, Jimenez-Movilla M, Dean J. FIGLA, a basic helix-loop-helix transcription factor, balances sexually dimorphic gene expression in postnatal oocytes. Mol Cell Biol. 2010 Jul;30(14):3661–71.

62. Raz E. The function and regulation of vasa-like genes in germ-cell development. Genome Biol. 2000 Sep 1;1(3):REVIEWS1017.

63. Campolo F, Gori M, Favaro R, Nicolis S, Pellegrini M, Botti F, et al. Essential role of Sox2 for the establishment and maintenance of the germ cell line. Stem Cells. 2013 Jul;31(7):1408–21.

64. Moore FL, Jaruzelska J, Fox MS, Urano J, Firpo MT, Turek PJ, et al. Human Pumilio-2 is expressed in embryonic stem cells and germ cells and interacts with DAZ (Deleted in AZoospermia) and DAZ-like proteins. Proc Natl Acad Sci U S A. 2003 Jan 21;100(2):538–43.

65. Nilsson EE, Doraiswamy V, Skinner MK. Transforming growth factor-beta isoform expression during bovine ovarian antral follicle development. Mol Reprod Dev. 2003 Nov;66(3):237–46.

66. Zarate-Garcia L, Lane SIR, Merriman JA, Jones KT. FACS-sorted putative oogonial stem cells from the ovary are neither DDX4-positive nor germ cells. Sci Rep. 2016 Jun 15;6(1):27991.

67. Huyhn K, Renfree MB, Graves JA, Pask AJ. ATRX has a critical and conserved role in mammalian sexual differentiation. BMC Dev Biol. 2011 Jun 14;11(1):39.

68. Yu C, Fan X, Sha QQ, Wang HH, Li BT, Dai XX, et al. CFP1 regulates histone H3K4 trimethylation and developmental potential in mouse oocytes. Cell Rep. 2017 Aug 1;20(5):1161–72.

69. Jiang Y, Zhang HY, Lin Z, Zhu YZ, Yu C, Sha QQ, et al. CXXC finger protein 1-mediated histone H3 lysine-4 trimethylation is essential for proper meiotic crossover formation in mice. Development. 2020 Mar 17;147(6):dev183764.

70. Moreno SG, Attali M, Allemand I, Messiaen S, Fouchet P, Coffigny H, et al. TGFbeta signaling in male germ cells regulates gonocyte quiescence and fertility in mice. Dev Biol. 2010 Jun 1;342(1):74–84.

71. Belli M, Shimasaki S. Molecular aspects and clinical relevance of GDF9 and BMP15 in ovarian function. Vitam Horm. 2018;107:317–48.

72. D’Aniello E, Paganos P, Anishchenko E, D’Aniello S, Arnone MI. Comparative Neurobiology of Biogenic Amines in Animal Models in Deuterostomes. Front Ecol Evol. 2020 Sep 25;8.

73. Bhat US, Shahi N, Surendran S, Babu K. Neuropeptides and behaviors: How small peptides regulate nervous system function and behavioral outputs. Front Mol Neurosci. 2021 Dec 2;14:786471.

74. Tian S, Zandawala M, Beets I, Baytemur E, Slade SE, Scrivens JH, et al. Urbilaterian origin of paralogous GnRH and corazonin neuropeptide signalling pathways. Sci Rep. 2016 Jun 28;6:28788.

75. Yañez-Guerra LA, Zhong X, Moghul I, Butts T, Zampronio CG, Jones AM, et al. Echinoderms provide missing link in the evolution of PrRP/sNPF-type neuropeptide signalling. Elife. 2020 Jun 24;9.

76. Escudero Castelán N, Semmens DC, Guerra LAY, Zandawala M, Dos Reis M, Slade SE, et al. Receptor deorphanization in an echinoderm reveals kisspeptin evolution and relationship with SALMFamide neuropeptides. BMC Biol. 2022 Aug 24;20(1):187.

77. Yañez-Guerra LA, Delroisse J, Barreiro-Iglesias A, Slade SE, Scrivens JH, Elphick MR. Discovery and functional characterisation of a luqin-type neuropeptide signalling system in a deuterostome. Sci Rep. 2018 May 8;8(1):7220.

78. Odekunle EA, Semmens DC, Martynyuk N, Tinoco AB, Garewal AK, Patel RR, et al. Ancient role of vasopressin/oxytocin-type neuropeptides as regulators of feeding revealed in an echinoderm. BMC Biol. 2019 Jul 31;17(1):60.

79. Semmens DC, Dane RE, Pancholi MR, Slade SE, Scrivens JH, Elphick MR. Discovery of a novel neurophysin-associated neuropeptide that triggers cardiac stomach contraction and retraction in starfish. J Exp Biol. 2013 Nov 1;216(Pt 21):4047–53.

80. Perillo M, Arnone MI. Characterization of insulin-like peptides (ILPs) in the sea urchin Strongylocentrotus purpuratus: insights on the evolution of the insulin family. Gen Comp Endocrinol. 2014 Sep 1;205:68–79.

81. Acevedo-Rodriguez A, Kauffman AS, Cherrington BD, Borges CS, Roepke TA, Laconi M. Emerging insights into hypothalamic-pituitary-gonadal axis regulation and interaction with stress signalling. J Neuroendocrinol. 2018 Oct;30(10):e12590.

82. Szukiewicz D. Current insights in prolactin signaling and ovulatory function. Int J Mol Sci. 2024 Feb 6;25(4):1976.

83. Clay CM, Cherrington BD, Navratil AM. Plasticity of anterior pituitary gonadotrope cells facilitates the pre-ovulatory LH surge. Front Endocrinol (Lausanne). 2020;11:616053.

84. Albertini D, Bromfield J. Soma-Germline Interactions in the Ovary: An Evolutionary Perspective. In: Oogenesis. Chichester, UK: John Wiley & Sons, Ltd; 2010. p. 103–13.

85. Wu Z, Yang L, He Q, Zhou S. Regulatory mechanisms of vitellogenesis in insects. Front Cell Dev Biol. 2020;8:593613.

86. Liu W, Xin Q, Wang X, Wang S, Wang H, Zhang W, et al. Estrogen receptors in granulosa cells govern meiotic resumption of pre-ovulatory oocytes in mammals. Cell Death Dis. 2017 Mar 9;8(3):e2662–e2662.

87. Planas JV, Athos J, Goetz FW, Swanson P. Regulation of ovarian steroidogenesis in vitro by follicle-stimulating hormone and luteinizing hormone during sexual maturation in Salmonid Fish1. Biol Reprod. 2000 May 1;62(5):1262–9.

88. Shirai H, Kanatani H, Taguchi S. 1-methyladenine biosynthesis in starfish ovary: action of gonad-stimulating hormone in methylation. Science. 1972 Mar 24;175(4028):1366–8.

89. Kanatani H, Shirai H, Nakanishi K, Kurokawa T. Isolation and identification of meiosis inducing substance in starfish Asterias amurensis. Nature. 1969 Jan;221(5177):273–4.

90. Johnson ET, Nicola T, Roarty K, Yoder BK, Haycraft CJ, Serra R. Role for primary cilia in the regulation of mouse ovarian function. Dev Dyn. 2008 Aug;237(8):2053–60.

91. Trombly DJ, Woodruff TK, Mayo KE. Roles for transforming growth factor beta superfamily proteins in early folliculogenesis. Semin Reprod Med. 2009 Jan;27(1):14–23.

92. Kipp JL, Golebiowski A, Rodriguez G, Demczuk M, Kilen SM, Mayo KE. Gene expression profiling reveals Cyp26b1 to be an activin regulated gene involved in ovarian granulosa cell proliferation. Endocrinology. 2011 Jan;152(1):303–12.

93. Ivarsson K, Sundfeldt K, Brännström M, Janson PO. Production of steroids by human ovarian surface epithelial cells in culture: possible role of progesterone as growth inhibitor. Gynecol Oncol. 2001 Jul;82(1):116–21.

94. Kloc M, Tworzydło W, Szklarzewicz T. Germline and somatic cell syncytia in insects. Results Probl Cell Differ. 2024;71:47–63.

95. Tingen C, Kim A, Woodruff TK. The primordial pool of follicles and nest breakdown in mammalian ovaries. Mol Hum Reprod. 2009 Dec;15(12):795–803.

96. Roco ÁS, Ruiz-García A, Bullejos M. Testis development and differentiation in amphibians. Genes (Basel). 2021 Apr 16;12(4):578.

97. Smaga CR, Bock SL, Johnson JM, Parrott BB. Sex determination and ovarian development in reptiles and amphibians: From genetic pathways to environmental influences. Sex Dev. 2023;17(2-3):99–119.

98. Hall GB, Long JA, Wood BJ, Bedecarrats GY. Germ cell dynamics during nest breakdown and formation of the primordial follicle pool in the domestic turkey (Meleagris gallopavo). Poult Sci. 2020 May;99(5):2746–56.

99. Spradling AC, Niu W, Yin Q, Pathak M, Maurya B. Conservation of oocyte development in germline cysts from Drosophila to mouse. Elife. 2022 Nov 29;11.

100. Wang Z, Liu CY, Zhao Y, Dean J. FIGLA, LHX8 and SOHLH1 transcription factor networks regulate mouse oocyte growth and differentiation. Nucleic Acids Res. 2020 Apr 17;48(7):3525–41.

101. De La Fuente R, Viveiros MM, Wigglesworth K, Eppig JJ. ATRX, a member of the SNF2 family of helicase/ATPases, is required for chromosome alignment and meiotic spindle organization in metaphase II stage mouse oocytes. Dev Biol. 2004 Aug 1;272(1):1–14.

102. Lesch BJ, Page DC. Genetics of germ cell development. Nat Rev Genet. 2012 Nov;13(11):781–94.

103. Nicholls PK, Schorle H, Naqvi S, Hu YC, Fan Y, Carmell MA, et al. Mammalian germ cells are determined after PGC colonization of the nascent gonad. Proc Natl Acad Sci U S A. 2019 Dec 17;116(51):25677–87.

104. Hickford DE, Frankenberg S, Pask AJ, Shaw G, Renfree MB. DDX4 (VASA) is conserved in germ cell development in marsupials and monotremes. Biol Reprod. 2011 Oct;85(4):733–43.

105. Hartung O, Forbes MM, Marlow FL. Erratum: Zebrafish vasa is required for germ-cell differentiation and maintenance. Mol Reprod Dev. 2016 Dec;83(12):1128–30.

106. Chen Y, Lin X, Dai J, Bai Y, Liu F, Luo D. Deletion of ddx4 ovary-specific transcript causes dysfunction of meiosis and derepress of DNA transposons in zebrafish ovaries. Biology (Basel). 2024 Dec 16;13(12).

107. Liu N, Han H, Lasko P. Vasa promotes *Drosophila* germline stem cell differentiation by activating *mei-P26* translation by directly interacting with a (U)-rich motif in its 3′ UTR. Genes Dev. 2009 Dec 1;23(23):2742–52.

108. Bristol-Gould SK, Kreeger PK, Selkirk CG, Kilen SM, Cook RW, Kipp JL, et al. Postnatal regulation of germ cells by activin: the establishment of the initial follicle pool. Dev Biol. 2006 Oct 1;298(1):132–48.

109. Iang Y. Hedgehog pathway inhibition causes primary follicle atresia and decreases female germline stem cell proliferation capacity or stemness. Stem Cell Res Ther. 2019;10.

110. Le Rolle M, Massa F, Siggers P, Turchi L, Loubat A, Koo BK, et al. Arrest of WNT/β-catenin signaling enables the transition from pluripotent to differentiated germ cells in mouse ovaries. Proc Natl Acad Sci U S A. 2021 Jul 27;118(30):e2023376118.

111. Zhang X, Gunewardena S, Wang N. Nutrient restriction synergizes with retinoic acid to induce mammalian meiotic initiation in vitro. Nat Commun. 2021 Mar 19;12(1):1758.

112. Moguilevsky JA, Wuttke W. Changes in the control of gonadotrophin secretion by neurotransmitters during sexual development in rats. Exp Clin Endocrinol Diabetes. 2001;109(4):188–95.

113. Liu X, Herbison AE. Dopamine regulation of gonadotropin-releasing hormone neuron excitability in male and female mice. Endocrinology. 2013 Jan;154(1):340–50.

114. Camille Melón L, Maguire J. GABAergic regulation of the HPA and HPG axes and the impact of stress on reproductive function. J Steroid Biochem Mol Biol. 2016 Jun;160:196–203.

115. Kaiser UB. Hyperprolactinemia and infertility: new insights. J Clin Invest. 2012 Oct;122(10):3467–8.

116. Xie Q, Kang Y, Zhang C, Xie Y, Wang C, Liu J, et al. The role of kisspeptin in the control of the hypothalamic-pituitary-gonadal axis and reproduction. Front Endocrinol (Lausanne). 2022 Jun 28;13:925206.

117. A.B. Chaet and R.A. McConnaughy. Physiologic Activity of Nerve Extracts. The Biological Bulletin. 1959;117:407–8.

118. Feng Y, Piñon Gonzalez VM, Lin M, Egertová M, Mita M, Elphick MR. Localization of relaxin-like gonad-stimulating peptide expression in starfish reveals the gonoducts as a source for its role as a regulator of spawning. J Comp Neurol. 2023 Sep;531(13):1299–316.

119. Chaiyamoon A, Tinikul R, Nontunha N, Chaichotranunt S, Poomtong T, Sobhon P, et al. Characterization of TRH/GnRH-like peptides in the sea cucumber, Holothuria scabra, and their effects on oocyte maturation. Aquaculture. 2020 Mar;518(734814):734814.

